## supplementary materials for "eQTL mapping using allele-specific gene expression"

#### A Data processing pipeline

Before going into the details of data processing steps for processing genotype and gene expression data from each dataset, we first illustrate a general pipeline (Figure 1). The steps of obtaining TReC per gene and per sample (the left part of the figure) and phased and imputed genotypes (the right side of the figure) are often standard steps in all the gene expression quantitative trait locus (eQTL) analyses. The middle section of the figure shows the steps of obtaining allele-specific reads at gene or SNP level, and it is an extra step to prepare data for eQTL mapping using allele-specific expression (ASE, or allele-specific read count, ASReC).

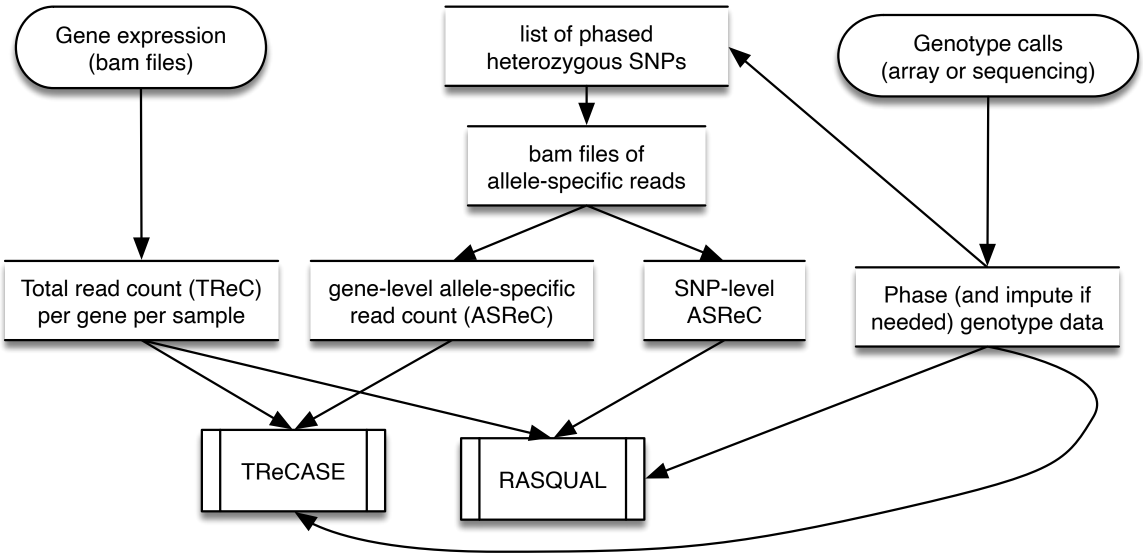

Figure 1: Data processing pipeline

##### A.1 Geuvadis dataset

The 280 samples of this eQTL dataset are lymphoblastoid cell lines that are part of the samples used in 1000 Genomes Project (1KGP) (1000 Genomes Project Consortium, 2015). The genotype data were obtained from SNP arrays and RNA-seq data were generated by the Geuvadis project (Lappalainen et al., 2013).

##### A.1.1 Genotype phasing and imputation

SNP genotype data were obtained from the same Geuvadis project. Using SHAPEIT2 (Delaneau et al., 2014) and IMPUTE2 (Howie et al., 2011, 2009), we phased and imputed genotypes in the following steps.

1. Convert unphased genotypes to PED/MAP format and ensure that genomic positions are converted to the genome reference that match the reference panel. This coordinate conversion can conveniently be done using liftover tool (Rhead et al., 2009).

2. Run SHAPEIT2 in check mode to get a list of mismatching SNPs. We have observed that there are a notable portion of SNPs labeled as strand mismatch after this step. To fix this problem we flip the SNPs with Plink (Purcell et al., 2007), repeat phasing in a check mode and compare resulting lists. If some of the SNPs present in first error list disappear in the second list they should be flipped and kept, and the rest of the mismatched SNPs can be supplied to SHAPEIT2 at the next step as an exclusion list.

3. Run SHAPEIT2 with exclusion list obtained in previous step.

4. Impute the pre-phased genotype data using IMPUTE2.

5. Using the imputed and phased data from the previous step, we obtain all heterozygous SNPs for each sample. Specifically, the main output file of imputation has 3 columns with genotype probabilities per SNP, which represent the probability of observing genotype $G = 0$, 1, or 2 respectively. We selected heterozygous locations (i.e., with high probability of $G = 1$) to output phased genotypes of these locations.

For both phasing and imputation, we used the 1000 Genome reference panel (Howie et al., 2011) (as of summer 2015) containing 2,504 samples with $\sim$82 million SNPs. Effective size of the population was set to the suggested value `--effective-size 20000` and random seed was set to 1234567. Imputation was done by splitting the genome into blocks of 5 Mb (no more than 7 Mb according to the instruction). We also used the same population size option as the one used in phasing step (`-Ne`

`20,000`), other options used inlcude `-align_by_maf_g` and `-seed 12345`.

The input data are genotypes of ∼2.2 million SNPs per sample for 2,123 samples of African or European descent. After phasing and imputation, we ended up with genotype data for ∼82 million SNPs per sample that include ∼2.2 million heterozygous SNPs per sample. Among all the imputed SNPs, around 6.5 million SNPs have MAF $> 0.05$ and were used as candidate eQTLs for eQTL mapping.

##### A.1.2 Processing RNA-seq

We downloaded raw RNA-seq data (in fastq format) of 462 samples that are part of the samples for the 1KGP (1000 Genomes Project Consortium, 2015). These RNA-seq data were generated by the Geuvadis consortium (Lappalainen et al., 2013), and it is available at `http://www.ebi.ac.uk/arrayexpress/experiments/E-GEUV-1/samples/`.

The RNA-samples were sequenced by the Illumina HiSeq2000 platform, with paired-end 75-bp reads. We mapped these RNA-seq reads to hg38 reference assembly using TopHat v2 (Trapnell et al., 2009), filtered with the following criteria: $\leq 3$ mismatches, read gap length $\leq 3$, read edit distance $\leq 3$, and read realign edit distance equals to 0.

We filtered RNA-seq reads with average base quality $\geq 30$, mapping quality $\geq 20$, and keeping only those uniquely mapped reads, by the `prepareBAM` function from R package `asSeq`. In a typical sample, most of the RNA-seq reads pass these filters.

For each sample, given the list of heterozygous SNPs obtained from phased and imputed genotype data, we filter out allele-specific reads using R function `extractASReads` from `asSeq` package (Sun, 2012). This step generates 3 bam files for each sample labeled with $hap1$, $hap2$ or $hapN$, which contain the RNA-seq reads that match haplotype 1, haplotype 2, or with conflicting information. Which haplotype is defined as haplotype 1 or haplotype 2 in each sample is arbitrary. The size of $hapN$ file should be much smaller than $hap1$ or $hap2$. Otherwise, there may be some systematic mismatch between RNA-seq bam files and the SNP list, e.g., they were generated using different human genome references. Apparent imbalance in *hap*1 and *hap*2 file sizes often suggest some problems in the data preparation steps.

Next we obtained the Total Read Count (TReC) and allele-specific read count (ASReC) per gene using R function `summarizeOverlaps` from `GenomicAlignments` package (Lawrence et al., 2013) using Gencode version 21 (GRCh38).We observed that for some samples, many genes have extreme proportions of reads attributed to one haplotype - this is likely due inconsistency between genotype data and RNA-seq data. We discarded such samples and used the remaining 280 samples for eQTL mapping. The total number of reads mapped to genes in these 280 samples vary from 16 to 82 million with fraction of allele-specific reads from 2.6% to 6.3% (Figure 2).

For RASQUAL we produced SNP level allele-specific counts using the same imputed SNPs and mapped bam files as input to `ASEReadCounter` from GATK package (McKenna et al., 2010).

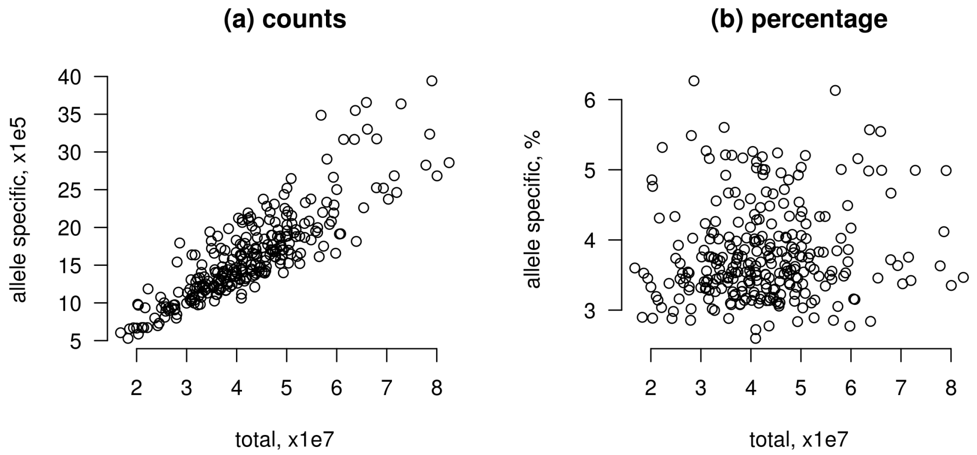

Figure 2: Summary of total read counts (x-axis) versus total number of allele-specific reads (a) or the percentage of reads being allele-specific (b) across all genes per sample for the Geuvadis data. Each point indicates one of the 280 samples.

#### A.2 GTEx v7 dataset

All the eQTL results reported in our paper are based on GTEx v8 data. Though when we first started this project, the v8 data were not available and we use the whole blood data from v7 (GTEx Consortium, 2017) to compare different computational methods. Here we briefly described how the v7 data were processed. We downloaded mapped RNA-seq data (in bam format) of 427 whole blood samples through NCBI Sequence Read Archive (SRA) after applying for data access through dbGaP (access id phs000424.v7.p2). The RNA-samples were sequenced by the Illumina HiSeq2000 platform with paired-end 76-bp reads and were mapped to human genome references hg19. We filtered RNA-seq reads with average base quality $\geq 20$, mapping quality $\geq 20$, and keeping only those uniquely mapped reads, by the `prepareBAM` function from R package `asSeq`.

We downloaded genotype calls (in VCF format) of 635 samples (release v7, hg19) from dbGaP. Indels and multi-allelic SNPs were removed using `bcftools` and variants with missing rate large than 10% were filtered out. We performed phasing on all 635 samples, with the same reference panel and command used when phasing Geuvadis data and used the resulting output file to create a list of heterozygous SNPs for each individual. Then we applied the same approaches as for Geuvadis data to collect Total Read Count (TReC) and Allele-Specific Read Count (ASReC) per gene and per sample for TReCASE and RASQUAL.

We filtered out genes whose 75 percentile of gene expression is less than 20 or maximum gene expression is more than 25 million. The resulting 16,675 genes were included in the analysis. The total number of reads mapped to genes in whole blood samples vary from 1 to 30 million with fraction of allele-specific reads from 2.3% to 11.73%, with one apparent outlier (Sample ID: YEC3 labeled in red in Figure 3), and it was removed in the following analysis.

Covariates data including genotyping principal components, age, and gender was downloaded from `https://gtexportal.org/home/datasets`. 354 samples, having RNA-seq data from whole blood, genotype data and covariates data, were included in the following eQTL mapping.

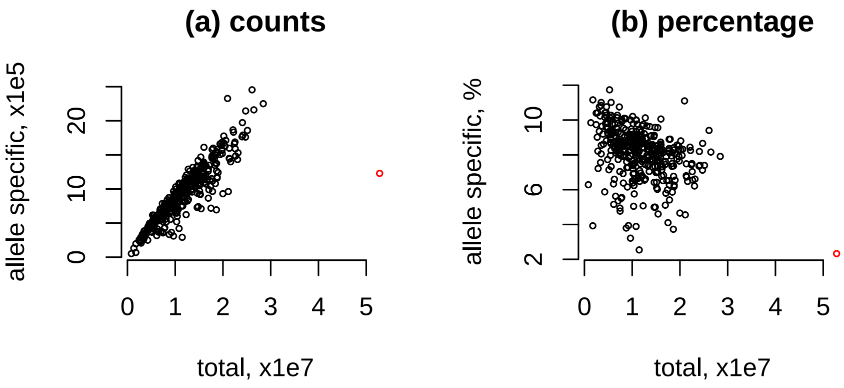

Figure 3: Summary of total read counts (x-axis) versus total number of allele-specific reads (panel (a)) or the percentage of reads being allele-specific (panel (b)) across all genes per sample for GTEx data. The red point indicates sample YEC3 has unexpected low proportion of allele-specific reads and it is excluded from further analysis.

#### A.3 GTEx v8 dataset

##### A.3.1 Genotype and covariates data

Using the GTEx v8 release, we conducted eQTL analyses for 22 GTEx tissues with sample size larger than 300 together with an additional artery tissue of artery-coronary, and 5 brain tissues: Caudate (basal ganglia), Cerebellar Hemisphere, Cortex, Frontal Cortex, and Nucleus accumbens (basal ganglia). We used the same set of covariates as GTEx Consortium et al. (2020) in eQTL analysis. The covariate data were downloaded from GTEx data portal: file `GTEx_Analysis_v8_eQTL_covariates.tar.gz` from `https://gtexportal.org/home/datasets`. The covariates include genotype principal components, PEER factors derived from gene expression data, and a few other variables such as sex, pcr, and platform. Following the GTEx study, we did not use age as a covariate for eQTL mapping, but we used age in dynamic eQTL studies. The age information were obtained from file

GTEx_Analysis_v8_Annotations_SubjectPhenotypesDS.txt on GTEx data portal.

The genotype calls from whole genome sequencing were part of the GTEx V8 protected data, that are available at https://anvil.terra.bio/#workspaces/anvil-datastorage/AnVIL_GTEx_V8_hg38. Access of such data require approved access to the GTEx data on dbGaP. The exact file we used is: GTEx_Analysis_2017-06-05_v8_WGS_VCF_files_GTEx_Analysis_2017-06-05_v8_WholeGenomeSeq_838Indiv_Analysis_Freeze.SHAPEIT2_phased.vcf. Because the genotype data were from whole genome sequencing, there is no need for further imputation, and it is already phased. Therefore, we just directly extract the list of heterozygous SNPs for each individual and use this list to calculate ASE.

##### A.3.2 Processing RNA-seq on a cloud computing platform

The Geuvadis dataset has manageable size so that it can be downloaded and processed in a local computational environment, either a desktop or a computer cluster. In GTEx analysis, we seek to reanalyze the data from 28 tissues. For the GTEx dataset and possibly many other datasets, the raw data are too large to download to process locally. We developed a pipeline for data processing using the data stored in cloud. Although this pipeline was designed for the GTEx data saved in NHGRI AnVIL https://anvil.terra.bio/, it can be applied to other datasets as well. This pipeline is available at a GitHub repository, gtex_AnVIL: https://github.com/Sun-lab/gtex_AnVIL.

This pipeline processes the RNA-seq data sample by sample. For each sample, the input are raw RNA-seq data (e.g., bam files), a list of phased heterozygous SNPs, and annotation of genes, e.g., the transcript/exon annotations from GECODE (Frankish et al., 2019). The output is a text file with 4 columns of counts that correspond to total counts and counts haplotype 1 (hap1), haplotype 2 (hap2), and haplotype N (hapN). The definitions of hap1, hap2, and hapN are the same as those defined for the Geuvadis dataset. There are also row names that correspond to gene symbol.

An example is shown below.

```
sample1.bam sample1_hap1.bam sample1_hap2.bam sample1_hapN.bam
ENSG00000000003.14 9 0 0 0
ENSG00000000005.5 0 0 0 0
ENSG00000000419.12 209 0 0 0
ENSG00000000457.13 303 51 40 0
ENSG00000000460.16 49 2 1 0
ENSG00000000938.12 29290 58 42 0
ENSG00000000971.15 46 1 8 0
ENSG00000001036.13 262 32 30 0
```

The pipeline is written by Workflow Description Language (WDL) (`collect_TReC_ASReC.wdl` in gtex_AnVIL repository), which calls computational engine embedded in a Docker file (folder `Docker` in gtex_AnVIL repository). An R script within the docker performed quality controls select RNA-seq reads with average base quality $\geq 20$, mapping quality $\geq 20$ and additional filtering provided by on R package `Rsamtools`:

```
flag1  = scanBamFlag(isUnmappedQuery=FALSE, isSecondaryAlignment=FALSE,
                     isDuplicate=FALSE, isNotPassingQualityControls=FALSE,
                     isSupplementaryAlignment=FALSE, isProperPair=TRUE)

param1 = ScanBamParam(flag=flag1, what="seq", mapqFilter=255)
```

### B Supplementary Methods

#### B.1 Probability distributions used by TReCASE and RASQUAL

Because we perform eQTL analysis for each gene separately, in the following section we define the probability distribution for one gene and omit the index for gene.

##### B.1.1 TReCASE definition

Let $T_i$ be the total read count (TReC) for a gene of interest in the $i$-th sample, with $i = 1, \ldots, n$. In TReCASE model, $T_i$ is assumed to follow a negative binomial distribution with density function defined as:

$$f_{NB}(T_i = t_i; \mu_i, \phi) = \frac{\Gamma(t_i + 1/\phi)}{\Gamma(t_i + 1)\Gamma(1/\phi)} \left(\frac{1/\phi}{1/\phi + \mu_i}\right)^{1/\phi} \left(\frac{\mu_i}{1/\phi + \mu_i}\right)^{t_i}, \tag{1}$$

where $\mu_i$ is sample-specific mean value and $\phi$ is an over-dispersion parameter. It can be show that the variance of $T_i$ is

$$\mathrm{Var}(T_i) = \mu_i + \mu_i^2 \phi, \tag{2}$$

and in the limiting case when $\phi \to 0$, the negative binomial distribution converges to a Poisson distribution.

Sample-specific mean value $\mu_i$ is defined as a function of some covariates:

$$\log(\mu_i) = \beta_0 + \beta_\kappa \log(\kappa_i) + \sum_{u=1}^{p} \beta_u c_{iu} + \eta_i, \tag{3}$$

where $\beta_0$ is an intercept, $\beta_\kappa$ is the coefficient for log-read-depth $\log(\kappa_i)$, $\beta_u$, $u = 1, ..., p$, is the regression coefficient for the $u$-th covariate (e.g., age, gender, batch effects etc.), and $\eta_i$ is the genetic effect. Given a candidate eQTL with two alleles $A$ and $B$, let $g_i$ be its genotype, defined as the number of $B$ alleles such that $g_i = 0$, 1, or 2 for genotype of homozygous $A$ allele, heterozygous, or homozygous $B$ allele, respectively. Then $\eta_i$ is:

$$\eta_i = \begin{cases} 0 & \text{if } g_i = 0 \\ \log[1 + \exp(b_0)] & \text{if } g_i = 1 \\ b_0 & \text{if } g_i = 2 \end{cases}, \tag{4}$$

where $b_0$ is the genetic effect, defined as log ratio of gene expression for genotype $BB$ vs. $AA$. The derivation of $\eta_i$ follows from the assumption that gene expression is additive across the two alleles before log-transformation, see equations (3)-(7) of Sun (2012) for details.

For TReCASE, allele-specific expression (ASE), or allele-specific read counts (ASReCs) are collected for two haplotypes at gene-level, whereas the ASReCs for RASQUAL are measured at SNP level, as described in the next section. For the $i$-th sample, denote the two ASReCs for haplotypes 1 and 2 by $N_{i1}$ and $N_{i2}$, so that total ASReC for the $i$-th sample is $N_i = N_{i1} + N_{i2}$. Note that for each sample, which haplotype is defined as haplotype 1 is arbitrarily decided. The distribution of $N_{i1}$ given $N_i$ can be modeled by a beta-binomial distribution:

$$f_{BB}(N_{i1} = n_{i1}; N_i = n_i, \alpha_i, \beta_i) = \binom{n_i}{n_{i1}} \frac{\Gamma(n_{i1} + \alpha_i)\Gamma(n_i - n_{i1} + \beta_i)}{\Gamma(n_i + \alpha_i + \beta_i)} \frac{\Gamma(\alpha_i + \beta_i)}{\Gamma(\alpha_i)\Gamma(\beta_i)}, \quad (5)$$

where $\alpha_i$ and $\beta_i$ are sample specific parameters and they are connected with expected proportion of reads of haplotype 1 (denoted by $\pi_i$) and over-dispersion (denoted by $\theta$) of this beta-binomial distribution by

$$\pi_i = \frac{\alpha_i}{\alpha_i + \beta_i} \text{ and } \theta = \frac{1}{\alpha_i + \beta_i}. \quad (6)$$

The variance of this beta-binomial distribution is

$$\text{Var}(N_{i1}) = n_i \pi_i (1 - \pi_i) \frac{1 + n_i \theta}{1 + \theta},$$

which converges to the variance of binomial distribution when $\theta = 0$.

In an alternative parametrization, the over-dispersion parameter can be rescaled to interval [0,1): $\rho = \theta/(1+\theta)$ (Paul et al., 2005). We will switch to this parametrization in section C.4.4 to study inflation introduced by splitting reads across several SNPs within a gene.

Let $G_i$ be the ordered genotype of the candidate eQTL, which takes values 0, 1, 2, and 3 for ordered genotype $AA$, $AB$, $BA$, and $BB$. An ordered genotype is defined based on the order of haplotype 1 followed by haplotype 2. For example, $AB$ indicates that $A$ is on haplotype 1 and $B$ is on haplotype 2. Given $G_i$, we can model $\pi_i$ as a function of genetic effect $b_0$:

$$\log\left(\frac{\pi_i}{1-\pi_i}\right) = \begin{cases} -b_0 & \text{if } G_i = AB \\ b_0 & \text{if } G_i = BA \\ 0 & \text{if } G_i = AA \text{ or } BB \end{cases}. \tag{7}$$

The genetic effect $b_0$ for TReC (equation (4)) and ASE (equation (7)) are the same and thus it can be estimated by combining the data from TReC and ASE.

##### B.1.2 RASQUAL definition

To illustrate the difference between TReCASE and RASQUAL, we only consider the basic RASQUAL model without additional features such as capturing sequencing/mapping error or reference allele mapping bias. In addition, we change the notations used by RASQUAL to be consistent with the notations of TReC to facilitate comparison. We will adopt two terms used in the RASQUAL paper: *cis*-regulatory SNP (**rSNP**) and feature SNP (**fSNP**). An rSNP is a candidate eQTL SNP and a fSNP is a SNP within exonic region of a gene where ASReC can be measured.

RASQUAL also models TReC $T_i$ by a negative binomial distribution. Though instead of a common over-dispersion parameter for all samples, it has a sample-specific over-dispersion parameter $\phi_i$:

$$f_{NB}(T_i = t_i; \mu_i, \phi_i) = \frac{\Gamma(t_i + 1/\phi_i)}{\Gamma(t_i+1)\Gamma(1/\phi_i)} \left(\frac{1/\phi_i}{1/\phi_i + \mu_i}\right)^{1/\phi_i} \left(\frac{\mu_i}{1/\phi_i + \mu_i}\right)^{t_i}. \tag{8}$$

The mean and over-dispersion is parametrized by

$$\mu_i = \lambda K_i Q_i \text{ and } \phi_i = \frac{1}{\theta_R K_i Q_i}, \tag{9}$$

where $\lambda$ is a scaling parameter for absolute mean of coverage depth of this gene, $K_i$ is a sample specific offset reflecting library size and other a priori estimated size factors, $Q_i$ is genetic effect defined later, and $\theta_R$ is a scaling parameter for over-dispersion.

Then the variance of $T_i$ is $\text{Var}(T_i) = \mu_i + \mu_i^2\phi_i = \mu_i(1 + \lambda/\theta_R)$.

Recall that $g_i$ denotes the genotype of a candidate eQTL, and it equals to 0, 1, or 2 for genotype $AA$, $AB/BA$, or $BB$. RASQUAL quantifies genotype effect by

$$Q_i = \begin{cases} 2(1-\pi) & \text{if } g_i = 0, \\ 1 & \text{if } g_i = 1, \\ 2\pi & \text{if } g_i = 2. \end{cases}$$

Then the sample-specific mean value in log scale is

$$\log(\mu_i) = \log(\lambda K_i Q_i) = \log(\lambda) + \log(K_i) + \log(Q_i). \quad (10)$$

Comparing the mean value of TReCASE (equation (3)) versus RASQUAL (equation (10)), it is easy to see that $\log(K_i)$ captures the effects of all the covariates. Although it is not explicitly mentioned in the RASQUAL paper, we expect the scale of $K_i$ to be around 1 because they assume $\lambda$ captures the absolute mean value $\mu_i$. We can also see the correspondence of genetic effect between TReCASE ($b_0$) and RASQUAL ($\pi$) is $\log[\pi/(1-\pi)] = b_0$.

While there is minor difference between TReCASE and RASQUAL in their TReC model, the major difference is their specifications for the ASE data. In RASQUAL, ASReCs are measured for each feature SNP (fSNP) (with SNP index $l = 1, \ldots, L$), denoted by $N_{il0}$ and $N_{il1}$ (with $N_{il} = N_{il0} + N_{il1}$). Then given $N_{il}$, $N_{il1}$ is modeled by a beta-binomial distribution

$$f_{BB}(N_{il1} = n_{il1}; N_{il} = n_{il}, \alpha_{il}, \beta_{il}) = \binom{n_{il}}{n_{il1}} \frac{\Gamma(n_{il} - n_{il1} + \alpha_{il})\Gamma(n_{il1} + \beta_{il})}{\Gamma(n_{il} + \alpha_{il} + \beta_{il})} \frac{\Gamma(\alpha_{il} + \beta_{il})}{\Gamma(\alpha_{il})\Gamma(\beta_{il})}. \quad (11)$$

Let $0 < h \leq 1/L$ be the relative proportion ASReC contributed by each fSNP, then

$$\alpha_{il} = h\theta_R K_i Q_{il0} \text{ and } \beta_{il} = h\theta_R K_i Q_{il1},$$

where $Q_{il0}$ and $Q_{il1}$ quantify the number of ASReC from haplotype 0 and haplotype 1, and they are defined in Table 6. Note that $Q_{il} = Q_{il0} + Q_{il1} = Q_i$, which only

Table 6: Relative mean for ASReCs. This table is taken from supplementary table 4 of Kumasaka et al. (2016). The first column defines the ordered genotype for rSNP (reference SNP or candidate eQTL) and fSNP (feature SNP where ASReC is measured) where 0 and 1 indicate reference and alternative allele, respectively.

| rSNP,fSNP | $Q_{il0}$ | $Q_{il1}$ | $Q_{il} = Q_i$ |
|---|---|---|---|
| 00,00 | $2(1-\pi)$ | 0 | $2(1-\pi)$ |
| 00,01 or 00,10 | $(1-\pi)$ | $(1-\pi)$ | $2(1-\pi)$ |
| 00,11 | 0 | $2(1-\pi)$ | $2(1-\pi)$ |
| 01,00 or 10,00 | 1 | 0 | 1 |
| 01,10 or 10,01 | $\pi$ | $(1-\pi)$ | 1 |
| 01,01 or 10,10 | $(1-\pi)$ | $\pi$ | 1 |
| 01,11 or 10,11 | 0 | 1 | 1 |
| 11,00 | $2\pi$ | 0 | $2\pi$ |
| 11,01 or 00,10 | $\pi$ | $\pi$ | $2\pi$ |
| 11,11 | 0 | $2\pi$ | $2\pi$ |

depends on the genotype of the *cis*-regulatory SNP (rSNP, or candidate eQTL) but not the genotype of the $l$-th fSNP.

Then the expected proportion of reads of haplotype 1 and over-dispersion (denoted by $\vartheta$) of this beta-binomial distribution are

$$\pi_i = \frac{\alpha_i}{\alpha_i + \beta_i} = \frac{Q_{il1}}{Q_i} \text{ and } \vartheta = \frac{1}{\alpha_i + \beta_i} = \frac{1}{h\theta_R K_i Q_i}. \tag{12}$$

The variance of this beta-binomial distribution is

$$\text{Var}(N_{il1}) = n_{il1}\pi_i(1-\pi_i)\frac{1 + n_i\vartheta}{1+\vartheta},$$

which converges to the variance of binomial distribution when $\vartheta = 0$.

In the RASQUAL paper, the authors further set $h = 1$ for the following reasons quoted from page 42 of their supplementary materials:

> "Here the constant $h$ reflecting the proportion of total AS count at each feature SNP is arbitrary ($0 < h \leq 1/L$ for $L > 0$; otherwise $h = 0$).

However, in our experience, $h < 1/L$ usually gives worse result in terms of power and fine-mapping than $h \approx 1$. This is partly because the larger the number of feature SNPs $L$ is, the smaller the proportion $h$ each feature SNP accounts for, resulting in overestimation of the dispersion parameter $\hat{\theta}$, resulting in more significant associations in hypothesis testing for features with larger $L$. To avoid this issue we set $h = 1$ to penalize the over-dispersion parameter more for large $L$."

Since $h$ denotes the proportion ASReC from each fSNP, it counter-intuitive to set it to be 1. It appears to be an ad-hoc solution to reduce the significance level, especially for the genes with large number of fSNPs. In fact, as shown in this paper, even when $h = 1$, RASQUAL still has inflated type I error, and the degree of inflation increases with the number of fSNPs (Figure 5B in main text). However, the reason is not overestimation of the dispersion parameter $\hat{\theta}$. The estimate of the dispersion parameter is often accurate. It is the mis-specified likelihood model that leads to underestimate of the variance of the eQTL effect size.

##### B.1.3 Definition of RASQUAL-like method: TReCASE-RL

To facilitate more pointed comparison between TReCASE and RASQUAL, in addition to running RASQUAL, we also implemented a modification of TReCASE to adopt two key assumptions made by RASQUAL but not by TReCASE: (1) Equating the over-dispersion parameters of the negative binomial distribution for TReC and the beta-binomial distribution for ASE; (2) Treating ASReC of each fSNP as independent beta-binomial observation. Specifically, for the $i$-th sample and the $l$-th SNP, we denote the two ASReCs for haplotypes 1 and 2 by $N_{il1}$ and $N_{il2}$. The distribution of $N_{il1}$ given $N_{il} = N_{il1} + N_{il2}$ is modeled by a beta-binomial distribution.

$$\begin{aligned} & f_{BB}(N_{il1} = n_{il1}; N_{il} = n_{il}, \alpha_{il}, \beta_{il}) \\ & \quad = \binom{n_{il}}{n_{il1}} \frac{\Gamma(n_{il1} + \alpha_{il})\Gamma(n_{il} - n_{il1} + \beta_{il})}{\Gamma(n_{il} + \alpha_{il} + \beta_{il})} \frac{\Gamma(\alpha_{il} + \beta_{il})}{\Gamma(\alpha_{il})\Gamma(\beta_{il})}, \end{aligned} \tag{13}$$

where $\alpha_{il}$ and $\beta_{il}$ are SNP specific parameters connected with expected proportion of reads of haplotype 1 (denoted by $\pi_i$) and over-dispersion (denoted by $\theta$) of this beta-binomial distribution by

$$\pi_{il} = \frac{\alpha_{il}}{\alpha_{il} + \beta_{il}} \text{ and } \theta = \frac{1}{\alpha_{il} + \beta_{il}}. \tag{14}$$

We call the modified TReCASE model as TReCASE-RL, where RL stands for "RASQUAL Like". By comparing TReCASE and TReCASE-RL, we can illustrate the consequence of these two assumptions.

###### B.1.4 The over-dispersion parameters of the three models

We summarize the over-dispersion parameters of three models: TReCASE, RASQUAL, and TReCASE-RL, in Table 7. The over-dispersion parameters of TReCASE are constants across all samples. In contrast, RASQUAL's over-dispersion parameters vary across samples because they depend on $K_i$ and $Q_i$. We expect both $K_i$ and $Q_i$ vary around the value of 1 and thus we may approximate the over-dispersion parameters of RASQUAL by assuming $K_i = Q_i = 1$. Then we can see the definition of over-dispersion parameters are in reverse scale. For TReCASE, larger over-dispersion parameter ($\phi$ or $\theta$) means larger over-dispersion, and for RASQUAL, larger over-dispersion parameter ($\theta_R$) means smaller over-dispersion. In our results, when we refer to an over-dispersion, it is the TReCASE over-dispersion.

Table 7: Over-dispersion parameters in TReCASE, RASQUAL, and TReCASE-RL models. RASQUAL model modifies the degree of over-dispersion variation with additional offset $K_i$ and relative genetic effect $Q_i$.

| Component | TReCASE | RASQUAL | TReCASE-RL |
|---|---|---|---|
| TReC | $\phi$ | $1/(\theta_R K_i Q_i) \approx 1/\theta_R$ | $\theta$ |
| ASE | $\theta$ | $1/(\theta_R K_i Q_i) \approx 1/\theta_R$ | $\theta$ |

##### B.2 Simulation setup

###### B.2.1 Simulation for TReC

We simulate total read count (TReC) using 4 covariates (including one that can be treated as library depth). Negative binomial over-dispersion parameter is set to be 0.01, 0.1 or 0.5. Genetic effect size $b_0$ is set to be 0, 0.125, 0.25, or 0.5. We simulate the TReC so that the median across samples is around 100.

##### B.2.2 Simulation for ASReCs without within-sample over-dispersion

In the basic simulation setup, we simulate ASReCs within a sample by a binomial distribution and introduce over-dispersion across samples. Specifically, we simulated data by the following steps:

1. We assume 10% of TReC are allele-specific (Figure 4(a)) and set the same genetic effect $b_0$ on ASReC as for TReC. The beta-binomial over-dispersion is defined as a fraction of negative binomial over-dispersion, with several values: 0, 1/8, 1/4, 1/2, 1 and 2 to cover the fractions observed in the Geuvadis data (Figure 4(b)). Fraction 0 corresponds to the extreme, but not unlikely scenario when ASReC follow a binomial distribution.

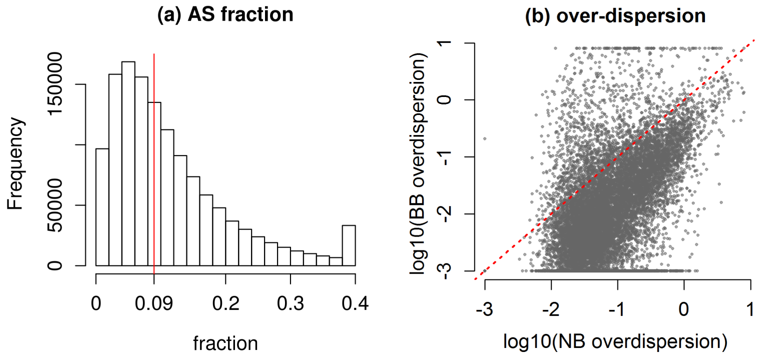

Figure 4: (a) The fraction of RNA-seq reads being allele-specific per gene per sample, given the ASReC is larger than 0, and truncated at 0.4. The vertical line indicates median. It is based on Geuvadis dataset of 280 samples. (b) The distribution of over-dispersion parameters estimated by TReCASE from the Geuvadis dataset of 280 samples. The BB over-dispersion is truncated at 0.001.

2. After calculating expected proportion of reads coming from allele B for each genotype $G$, denoted by $\pi_G$ we generate sample level expected proportions $\pi_i$ by sampling from a beta distribution with parameters $\alpha$ and $\beta$ such that $\pi_G = \alpha/(\alpha + \beta)$, and $\theta = 1/(\alpha + \beta)$.

3. Given $\pi_i$, we generate ASReC from a binomial distribution.

To simulate SNP-level ASReC, we modify step 3 as follows:

3a Uniformly distribute allele-specific reads (that can belong to either haplotype) among 2, 4, or 8 SNPs, and then simulate SNP-level ASReC on one haplotype by a binomial distribution using the same sample level proportion $\pi_i$.

##### B.2.3 Add within-sample over-dispersion for ASReC data

In order to simulate SNP level ASReC with over-dispersion across multiple SNPs within a sample, we modify step 3a as follows:

3b Uniformly distribute the allele-specific reads (that can belong to either haplotype) among 2, 4 or 8 SNPs, and then simulate SNP-level ASReC for each haplotype by a beta-binomial distribution with mean value $\pi_i$ and an over-dispersion parameter that equals to the between-sample beta-binomial over-dispersion.

This modification allows us to obtain ASReCs with over-dispersion within a sample and at the same time with more similarity within sample than between samples. Figure 5(c) in the main text use the results from this simulation setup.

##### B.2.4 To simulate RASQUAL style ASReC data

RASQUAL style SNP level ASReC data have the same over-dispersion across any two SNPs, either two SNPs within a sample or between samples. In other words, there is no extra over-dispersion across samples. To simulate such data, we modify step 2 to set the between-sample over-dispersion to be 0. After that we generate all SNP level ASReC from a beta-binomial distribution according to step 3b. This is equivalent to generating SNP level proportions $\pi_{il}$ from sample level proportions $\pi_i$ with the same over-dispersion $\theta$ without discrimination for the SNPs within and between samples.

##### B.2.5 Simulating the data with genotyping errors

To simulate genotyping errors, we randomly flip a fraction of genotypes from homozygous to heterozygous and from heterozygous to homozygous (fractions 0.05, 0.10 and 0.20 were used). The wrong genotypes have the following consequences.

1. TReCASE only uses heterozygous SNPs to collect ASReCs. In contrast, RASQUAL produces counts for homozygous SNPs too. Consequently, whenever we flip a heterozygous SNP to be homozygous, the read counts of this SNP are ignored by TReCASE, while still used by RASQUAL - it would have read counts of both alleles counted, but attach them to an incorrect homozygous genotype status.

2. When a truly homozygous SNP is flipped to be heterozygous, both methods still use such data. They both assume all the reads are from one of the two alleles.

### C Supplementary Results

#### C.1 Estimation of permutation p-value estimation using geoP

##### C.1.1 Evaluation of geoP using Geuvadis dataset

As a quick alternative to estimate permutation p-values, one might simply use eigenMT (Ongen et al., 2016) to estimate the effective number of tests, and then obtain the permutation p-value estimate by multiplying minimum p-value with the effective number of tests and truncating at 1. We evaluate this approach and our methods when estimating the relation between permutation p-value and nominal p-value using a linear regression or a logistic regression. As a gold standard to compare with, we estimated permutation p-values estimated by 10,000 permutations for 14,566 genes.

EigenMT tends to be conservative with a large number of false negatives (permutation p-value estimated using EigenMT is larger than a threshold $\alpha$ while the estimates by 10,000 permutations is smaller than $\alpha$), especially at less significant p-values (Table 8). This suggests that the eigenMT estimates of the number of tests is too large, particularly so for larger p-values. Overall the results based on linear fit is much more accurate than eigenMT (in terms of smaller number of false positives + false negatives), though it produces unbalanced false positives and false negatives, with more false positives at larger p-value cutoff and more false negatives at smaller p-value cutoffs (Table 8). Finally, the logistic regression has the the most accurate estimates with balanced numbers of false positives and false negatives (Table 8).

In terms of false classifications, we do not get as much improvement by using more than 100 grid points, particularly for logistic regression (glm) approach (Table 9). The performance of linear regression and logistic regression become similar at larger grid points and more significant p-values. Consequently, we suggest using 100 grid points to estimate permutation p-values, though even with 25 grid points we observe large improvement against eigenMT.

The estimation bias by eigenMT or linear model, as well as the advantage of using larger number of grid points, can be visualized by the scatter plots of permutation p-

Table 8: Permutation p-value misclassification by method. Misclassification by three methods for 14,566 genes from Geuvadis dataset. Quantifying by false positives (f.pos), false negatives (f.neg) and overall number of mis-classification (f.pos + f.neg). Here a positive outcome means the permutation p-value estimate is smaller than the cutoff value $\alpha$.

| | eigenMT | | | lm | | | glm | | | total |
|---|---|---|---|---|---|---|---|---|---|---|
| $\alpha$ | f.neg | f.pos | all | f.neg | f.pos | all | f.neg | f.pos | all | n.pos |
| 1e-1 | 1088 | 5 | 1193 | 7 | 97 | 104 | 27 | 18 | 45 | 6207 |
| 5e-2 | 694 | 4 | 871 | 5 | 85 | 90 | 34 | 25 | 59 | 5143 |
| 1e-2 | 275 | 10 | 285 | 21 | 36 | 57 | 17 | 30 | 47 | 3678 |
| 5e-3 | 219 | 12 | 207 | 31 | 24 | 55 | 17 | 30 | 47 | 3301 |
| 1e-3 | 85 | 25 | 110 | 52 | 26 | 78 | 28 | 39 | 67 | 2664 |

Table 9: Number of misclassifications of permutation p-value estimates for 25, 50, 100 or 200 grid points.

| permutation | eigenMT | lm | | | | glm | | | |
|---|---|---|---|---|---|---|---|---|---|
| p-value cutoff | | 25 | 50 | 100 | 200 | 25 | 50 | 100 | 200 |
| 0.1 | 1093 | 139 | 103 | 104 | 94 | 66 | 61 | 45 | 42 |
| 0.05 | 698 | 112 | 99 | 90 | 88 | 74 | 53 | 59 | 55 |
| 0.01 | 285 | 65 | 65 | 57 | 57 | 47 | 52 | 47 | 47 |
| 0.05 | 219 | 77 | 59 | 55 | 48 | 59 | 54 | 47 | 51 |
| 0.001 | 110 | 81 | 79 | 78 | 69 | 72 | 59 | 67 | 60 |

value estimates versus the "true values" estimated by 10,000 permutations (Figure 5).

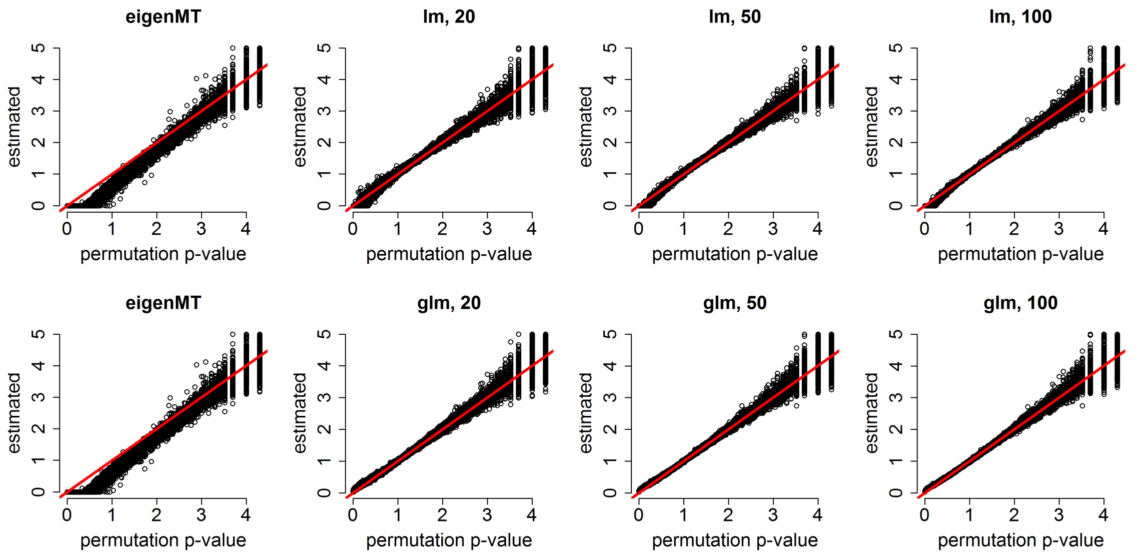

Figure 5: Permutation p-value estimation using three methods for Geuvadis dataset: eigenMT, linear model (lm), and logistic model (glm). The number following lm or glm is the number of grid points used. The x-axis are permutation p-values estimated by 10,000 permutations.

##### C.1.2 Evaluation of geoP using GTEx v7 dataset

Next we carried out similar evaluations of permutation p-value estimation using GTEx v7 data from whole blood. The conclusion is the same as those from Geuvadis dataset: glm has better performance than lm, which is better than eigenMT (Table 10) and 100 grid points is a good balance for computational cost and accuracy (Table 11). Finally, the scatter plot of permutation p-value estimates versus "true values" estimated by 10,000 permutations illustrate that glm with 100 grid points has the best estimation accuracy among all the methods (Figure 6).

Table 10: Permutation p-value misclassification by method.

Misclassification by three methods for 16,637 genes using GTEx v7 whole blood data. Quantifying by false positives, false negatives and overall number of mis-classification

| | eigenMT | | | lm | | | glm | | | total |
|---|---|---|---|---|---|---|---|---|---|---|
| $\alpha$ | f.neg | f.pos | all | f.neg | f.pos | all | f.neg | f.pos | all | n.pos |
| 1e-1 | 1000 | 0 | 1000 | 11 | 115 | 126 | 33 | 24 | 57 | 5001 |
| 5e-2 | 595 | 0 | 595 | 5 | 92 | 97 | 17 | 28 | 45 | 3887 |
| 1e-2 | 204 | 0 | 204 | 20 | 21 | 41 | 18 | 24 | 42 | 2489 |
| 5e-3 | 131 | 3 | 134 | 24 | 17 | 41 | 18 | 27 | 45 | 2177 |
| 1e-3 | 54 | 11 | 65 | 44 | 21 | 65 | 29 | 29 | 58 | 1708 |

Table 11: Number of misclassifications of permutation p-value estimates for 25, 50, 100 and 200 point grid.

| permutation | eigenMT | lm | | | | glm | | | |
|---|---|---|---|---|---|---|---|---|---|
| p-value cutoff | | 25 | 50 | 100 | 200 | 25 | 50 | 100 | 200 |
| 0.1 | 1000 | 184 | 121 | 126 | 106 | 82 | 67 | 57 | 60 |
| 0.05 | 595 | 100 | 108 | 97 | 89 | 55 | 43 | 45 | 44 |
| 0.01 | 204 | 52 | 50 | 41 | 38 | 48 | 45 | 42 | 34 |
| 0.005 | 134 | 65 | 60 | 41 | 48 | 50 | 45 | 45 | 46 |
| 0.001 | 65 | 71 | 62 | 65 | 57 | 55 | 56 | 58 | 54 |

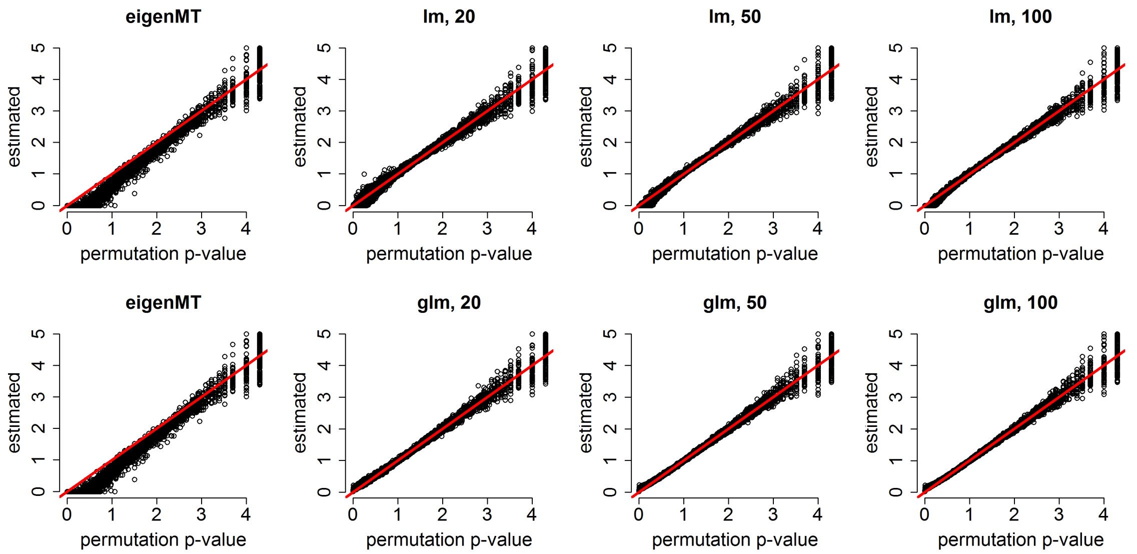

Figure 6: Permutation p-value estimation using three methods: eigenMT, linear regression and logistic regression using GTEx v7 whole blood dataset. The x-axis are permutation p-values estimated by 10,000 permutations.

#### C.2 Supplementary results for dynamic eQTLs.

##### C.2.1 Cell type proportion estimate from whole blood

In order to estimate cell type proportions, we need to know the relevant cell types of a tissue as well as cell type-specific gene expression. We choose to estimate cell type proportions in whole blood because there is a popular reference of 22 cell types for whole blood, known as LM22, which was compiled by Newman et al. (Newman et al., 2015). We first transformed the gene expression data from GTEx (v8, whole blood) to transcript per million (TPM), and then estimated cell type proportions using LM22 reference and CIBERSORTx web interface (`https://cibersortx.stanford.edu/`) (Newman et al., 2019).

##### C.2.2 Dynamic eQTLs related with CTCF and TP53

To evaluate the association between CTCF and TP53's expression and six potential confounders: top 4 PEER factors plus 2 genotype PCs, we fit a linear model with read-depth corrected CTCF or TP53's expression as response variable and the six potential confounders as covariates. These linear models can explain a substantial proportion of the variation of CTCF and TP53's expression (Figure 7a-b). See `https://github.com/Sun-lab/asSeq/blob/master/pipeline_GTEx/v8/cell_type_composition/step3_age_CTCF_TP53.pdf` for more details. After accounting for these six potential confounders, we can still identify some dynamic eQTLs and Figure 7c-d give two examples.

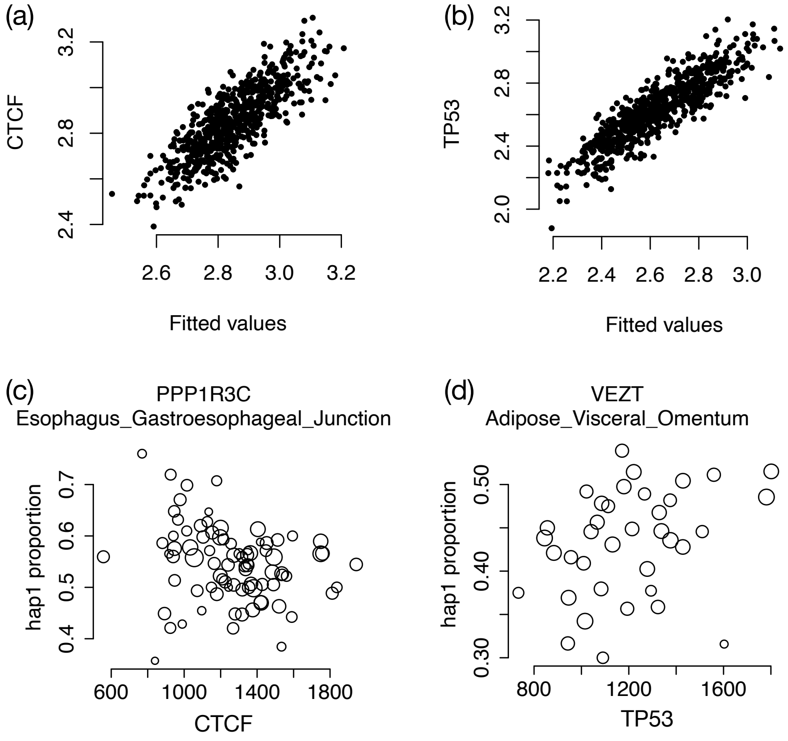

Figure 7: (a-b) Fitted values from a linear model with seven covariates (top 5 PEER factors plus 2 genotype PCs,) versus observed gene expression for *CTCF* and *TP53*. (c-d) Examples of dynamic eQTLs mediated by the expression of *CTCF* and *TP53*, respectively.

#### C.3 Evaluation of the binomial distribution assumption for ASReCs across multiple SNPs within one gene and one sample

TReCASE assumes the summation of ASReCs across multiple SNPs within a gene follows a beta-binomial distribution across samples. This assumption implies that the ASReCs across multiple SNPs within a gene and a sample should follow the same binomial distribution. If these SNP-level ASReCs in fact follow a beta-binomial distribution, their summation will not follow a beta-binomial distribution, though it may be approximated by a beta-binomial distribution. Here we evaluate this binomial distribution assumption. It cannot be evaluated for most (gene, sample) pairs because most such cases only have a few heterozygous SNPs with enough coverage. We used RNA-seq data from 30 HapMap trios generated from our earlier study (NCBI BioProject access number: PRJNA385599) (Zhabotynsky et al., 2019) to select a set of genes with allele-specific counts distributed across multiple SNPs. This dataset was used since it has higher read depth. We selected those (gene, sample) pairs such that the gene in the sample has at least 6 heterozygous SNPs, with at least 5 overlapping reads per SNP. We ended up with 4,005 (gene, sample) pairs matching these criteria, accounting for less than 1% of all (gene, sample) pairs.

For these 4,005 cases we tested how often the binomial assumption is violated. Deviation from such assumption can be tested using a score statistic developed by Tarone (1979). Since we do not have many SNPs, normal approximation of score statistic cannot be applied. Instead, we generated the null distribution of such score statistic using parametric bootstrap. For each gene, we estimated the proportion of reads from one allele, simulated ASReC for each SNP from a binomial distribution with this probability and the observed coverage and recalculated the score statistic. Then a p-value can be calculated as the relative frequency when the observed score statistic is more extreme than the ones from parametric bootstraps. Under the null we expect a uniform distribution of p-values, however we see an excess

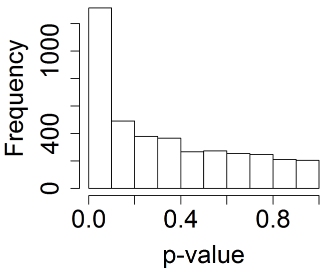

Figure 8: The distribution of p-values for testing deviation from binomial distribution across multiple SNPs of the same gene and within the same sample

of p-values in the category less than 0.1, which transforms to an estimate that the p-value is not uniform for ∼23% of the cases (Figure 8). Note that this 23% is among those selected 1% of the cases with enough heterozygous SNPs covered by RNA-seq reads, and thus only represent a very small proportion of all the data points in real data analysis.

#### C.4 Simulation Results

##### C.4.1 Simulation results under RASQUAL assumption

Under RASQUAL's assumption, the SNP-level ASReC follows a beta-binomial distribution, such that the similarity of the SNP-level ASReCs are the same for two SNPs within one sample versus two SNPs of two different samples. This is the simulation scenario described in Section B.2.4. It worth noting that this scenario is not supported by real data. As shown in Figure 8, when considering a subset of (gene, sample) pairs with at least 6 heterozygous SNPs and at least 5 overlapping reads per SNP, in 77% of the cases the ASReCs within a sample follow a binomial distribution. We mainly want to use this scenario to demonstrate that RASQUAL does control type I error if its model assumption is correct. In addition, TReCASE model is mis-specified in this scenario, and we demonstrate that TReCASE still has reasonable performance despite its model mis-specification.

As shown in Figure 9, both TReCASE and TReCASE-RL control type I error well except that TReCASE-RL is slightly conservative when negative binomial over-dispersion is 2 and beta-binomial over-dispersion is 0.25. The power of the two methods are similar. TReCASE has slightly higher power when the over-dispersion for beta-binomial distribution is small, and TReCASE-RL has slightly higher power when the over-dispersion for beta-binomial is large.

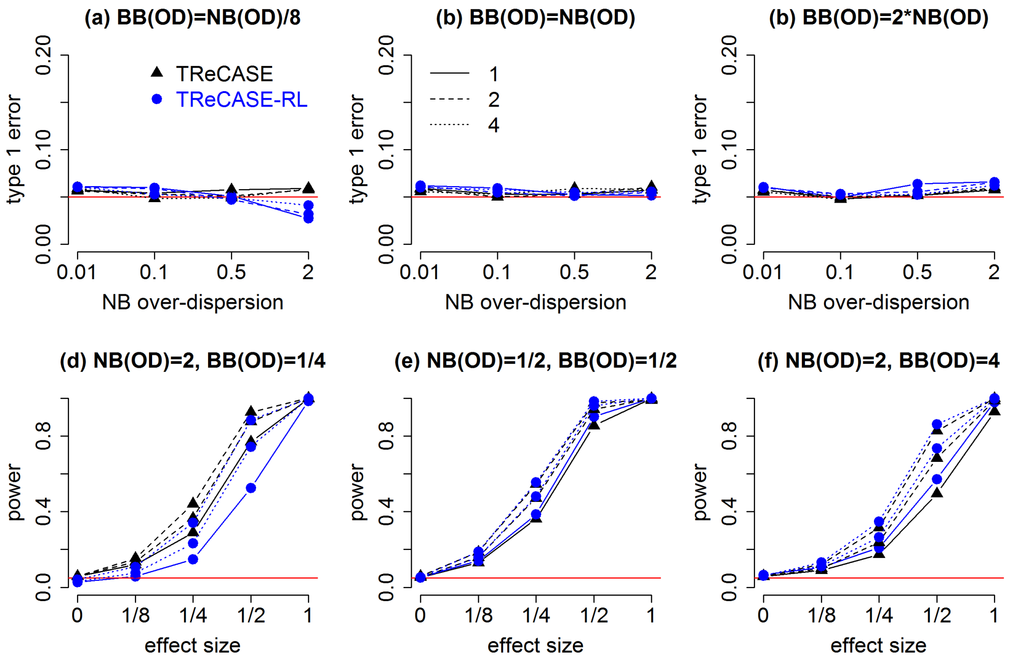

Figure 9: Evaluating TReCASE and TReCASE-RL using data simulated under RASQUAL assumption such that within sample over-dispersion is the same as between sample over-dispersion. Panels (a)-(c) present type I error and panels (d)-(f) present power. 10,000 genes were simulated for each effect size and over-dispersion profile. The three line types refer to the number of fSNPs per gene. We use sample size 64 in our illustrations.

##### C.4.2 Simulations results with within sample over-dispersion and additional between sample over-dispersion for ASReCs

This is the simulation setting where both models are mis-specified. Following the simulation described in Section C.3, we simulated ASReC data with within sample over-dispersion and extra between-sample over-dispersion. When simulating two SNPs per gene, TReCASE still manages to control type I error, while the RASQUAL style approach has inflated type I error and the inflation increases for larger beta-binomial over-dispersion (Figure 5(C) in main text). In the case of 4 or 8 SNPs per gene, the results are similar. TReCASE still controls type I error, and RASQUAL style approach shows even higher inflation of type I error (results not shown).

##### C.4.3 Evaluating models for the data simulated under TReCASE assumption

In previous section we have shown that for the majority of the genes the ASReC do not have within sample over-dispersion. This is a TReCASE style assumption. In this section, we simulated ASReC without within sample over-dispersion, and either combined them into one count or split them across two or four SNPs. Our simulation results show that in this scenario, TReCASE controls type I error reasonably well (Figure 10 (a)-(c)), while RASQUAL can either produce deflated type I error (Figure 10 (a)) or inflated type I error (Figure 10 (b)-(c)).

We also evaluated the effect of double counting by randomly selecting 10% of the reads from each SNP and adding them to a neighboring SNP. Double-counting further inflates type I errors in all three simulation settings (Figure 10 (d)-(f)).

Comparing the results between TReCASE-RL and RASQUAL, we concluded that RASQUAL is has some additional reasons to produces type I error inflation, not explained by equating total read counts and allele-specific counts over-dispersion or treating each SNP count as independent Beta-Binomial as implemented in TReCASE-RL version (Figure 10 (h-j)).

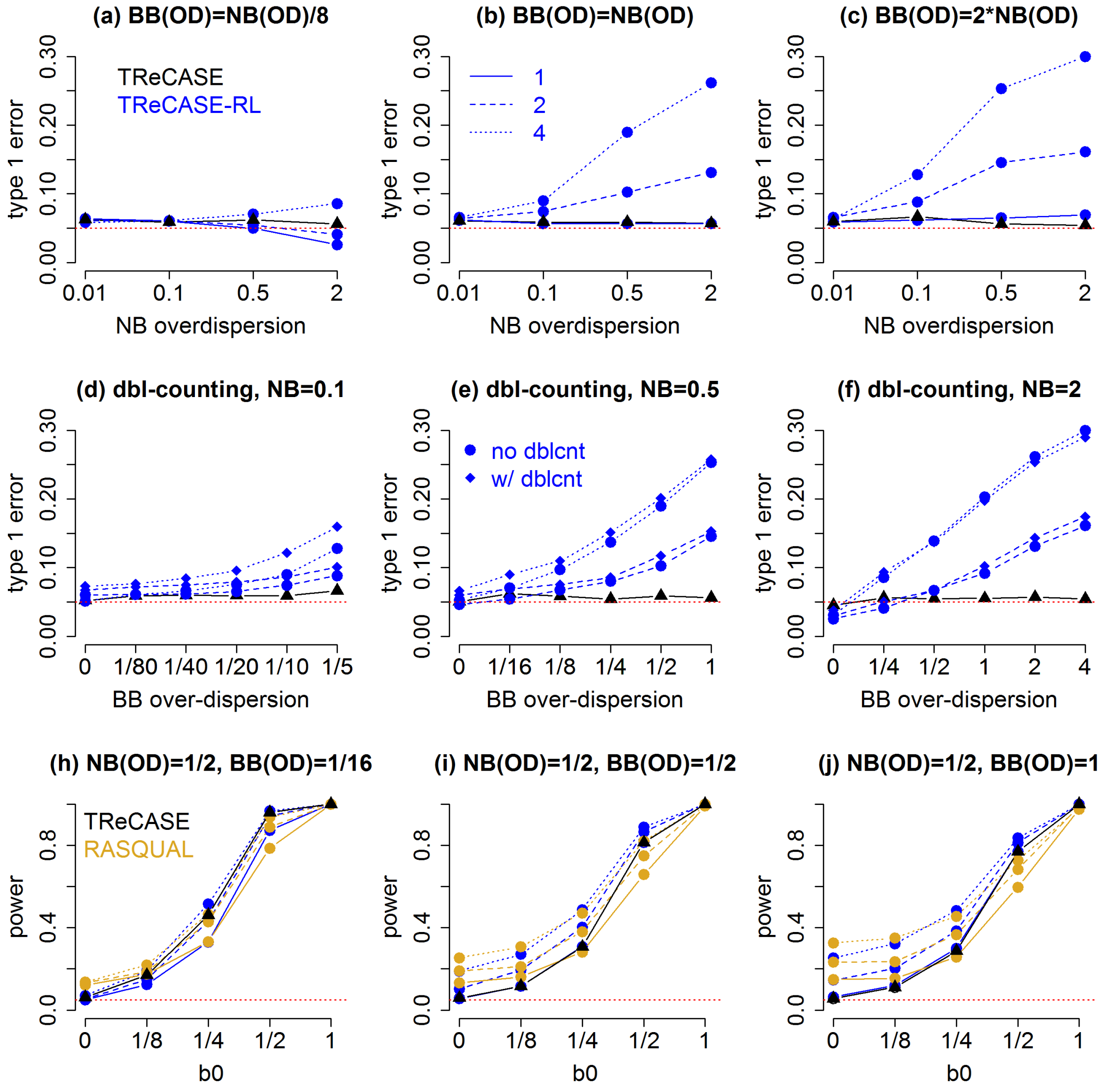

Figure 10: Evaluating TReCASE, TReCASE-RL, and RASQUAL models for the data simulated using TReCASE style assumptions. a)-(c) Evaluation of Type I error across different values of over-dispersion parameters. (d)-(f) Evaluation of type I error given double counting. (h)-(j) Evaluation of power and type I error across eQTL effect sizes. Results are presented for sample size 64.

##### C.4.4 Evaluation of the inflation of type I error by RASQUAL

In the previous section, we showed applying RASQUAL or TReCASE-RL on simulated RNA-seq data (using both TReC and ASReC), there is inflated type I error. In this section, we study this issue by ignoring TReC and concentrating on ASReCs without within-sample over-dispersion. We set beta-binomial over-dispersion to 0.1 or 0.5 and generate data under null hypothesis of no eQTL effect (proportion of either allele is set to be 0.5). We simulated data for 1,000 genes. For each gene, we first simulated the data assuming there is only one fSNP. Then we split the ASReCs uniformly to multiple fSNPs, while the total number of allele-specific reads are the same. For example, if there are $k$ reads per SNP for 8 SNPs, then there are $2k$ reads per SNP for 4 SNPs. We applied TReCASE-RL on these data. As shown in previous results, there is no inflation of type I error in one SNP scenario. In two SNP scenario, as expected, both likelihood ratio test-statistics (LRT) and -log(p-value) become larger than one SNP scenario (Figure 11).

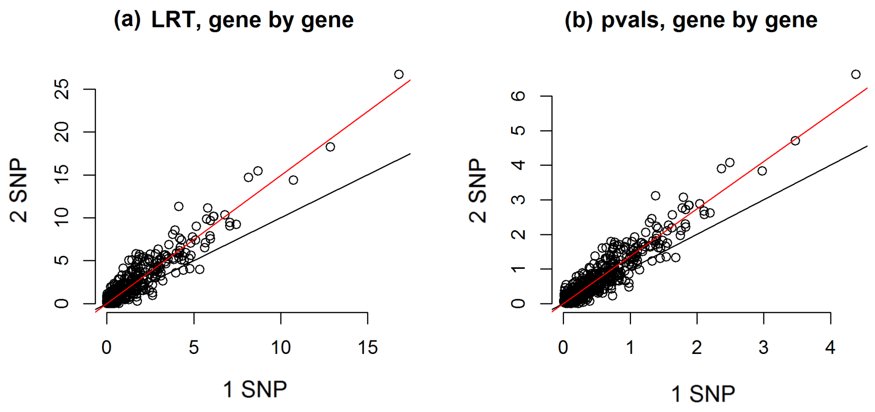

Figure 11: Illustration of the type I error inflation by TReCASE-RL. The data were simulated under null. Likelihood ratio test-statistics (LRT) and -log10(p-values) are compared for two situations: one fSNP or two fSNPs. Each point corresponds to one of 1000 genes. We observe that for the same gene once reads are split into two SNPs we tend to get more significant results.

To understand the underlying causes of type I error inflation, we examine the estimation of eQTL effect and over-dispersion across simulate replicates. We found that after splitting the allele-specific reads to 2, 4 or 8 fSNPs, the estimation of eQTL effects and over-dispersion are both unbiased (Figure 12). As the number of fSNPs increases, the variation of eQTL effect estimates remain similar while the variation of over-dispersion estimates increases. However, due to the mis-specification of likelihood model by RASQUAL, the mode-based standard deviation (sd:mod in Figure 12) estimates are smaller than observed sds (sd:obs in Figure 12) for both eQTL effects and over-dispersion. The under-estimation of sd for eQTL effects explains the inflation of type I error when we test for eQTL effect. Note that model-based sds do match well with empirical sds when there is only one fSNP. This is expected since there is no model mis-specification when there is only one fSNP.

We derive sd using the Fisher's information matrix derived by Paul et al. (2005), and to be consistent with their work, we quantify over-dispersion by $\rho = \theta/(1 + \theta)$, where $\theta$ was the over-dispersion parameter defined in Equation (1). To make the notation clear, we use legend OD(theta) or OD(rho) in the plots to indicate over-dispersion quantified by $\theta$ and $\rho$, respectively.

Next, we examine how the model-based sd estimates vary with respect to the number of allele-specific reads. As expected the sd estimates of either eQTL effect and over-dispersion decreases as ASReC increases (Figure 13). For eQTL effect size, we have observed in Figure 12 that the true sds are similar when the number of fSNPs is 1, 2, 4, or 8. Since the sd estimate of 1 SNP scenario is unbiased, the difference of the sd estimates with 1 SNP versus 2, 4, or 8 SNPs reflects bias of sd estimates due to model mis-specification (Figure 13 (a) and (c)). The standard deviation of over-dispersion parameter is of less interest in this study. Though we can see that when the number of ASReC is relatively large, the sds of over-dispersion tend to be under-estimated (Figure 13 (b) and (d)).

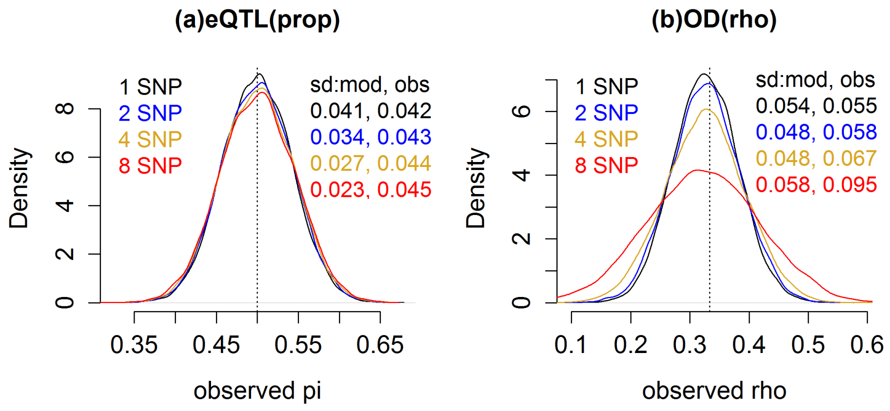

Figure 12: Distribution of eQTL effect and over-dispersion estimates. (a) Distribution of eQTL effect estimates in terms of $\pi$, the proportion of ASReC from one haplotype. (b) Distribution of over-dispersion estimates in terms of $\rho$, which is the rescaling of over-dispersion parameter $\theta$ to [0, 1) range by $\rho = \theta/(1+\theta)$. In the upper-right corner of each figure, we also list the model-based standard deviation estimate (sd:mod) using Fisher's information matrix (Paul et al., 2005) and empirical standard deviation estimate (sd:obs) across simulation replicates. The data were simulated under null of no genetic effect, and ASReCs were split into 1, 2, 4, and 8 SNPs. Simulation was done for sample size 64 and on average 10 allele-specific reads per sample.

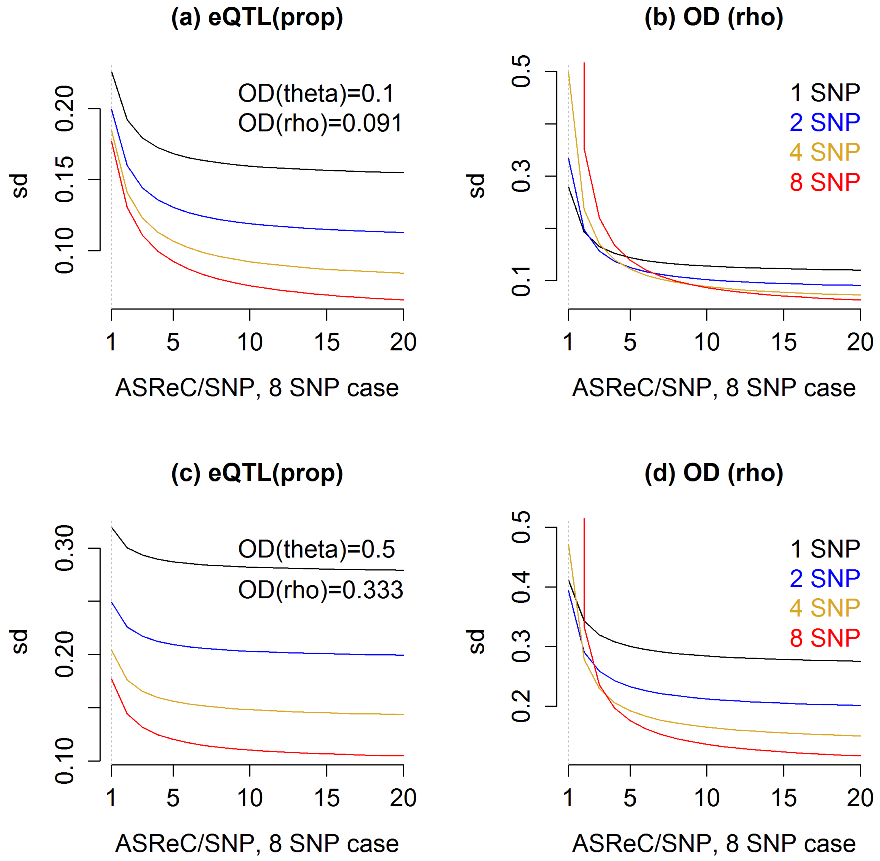

Figure 13: Model-based sd estimates by Fisher's information matrix under null. The sd estimates are evaluated for 1, 2, 4, and 8 fSNPs per gene. X-axis is the number of allele-specific reads for each of 8 fSNPs. For example, $x = 5$ means there are 5 reads for each of the 8 SNPs, or 10 reads for each of the 4 SNPs, or 20 reads for each of the 2 SNPs, or 40 reads for one SNP. Panels (a)-(b) present the simulation results for $\phi = 0.1$ and panels (c)-(d) present the simulation results for $\phi = 0.5$

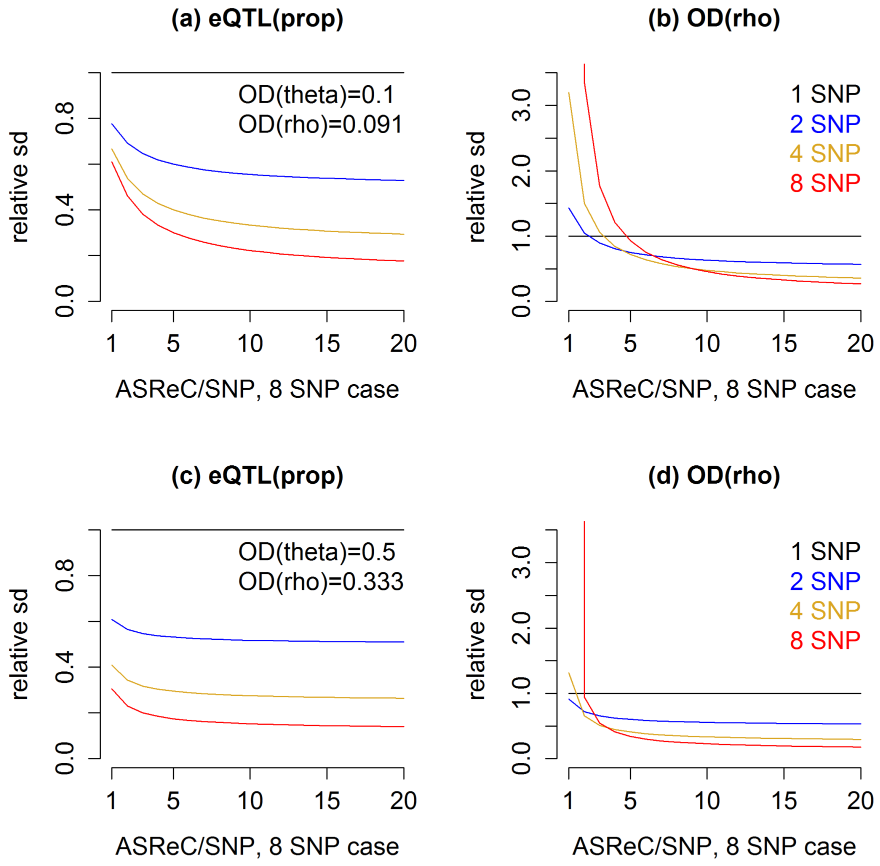

Figure 14: Model-based relative sd estimates under null.

The same as Figure 13, except that the y-axis is the relative sd estimates with respect to the sd estimate for 1 fSNP scenario.

##### C.4.5 Compare TReCASE versus RASQUAL when there are genotyping errors

Finally, we evaluate the results of TReCASE and RASQUAL when there are genotyping errors using simulated data. We simulated ASReC data without within sample over-dispersion (Section B.2.2), and then introduced genotyping errors as described in Section B.2.5, where we flipped certain fraction of genotypes from homozygous to heterozygous and from heterozygous to homozygous. When truly heterozygous SNPs are listed as homozygous, they will be discarded by TReCASE but used by RASQUAL, and the latter has a mechanism of correcting the genotype status if it encounters a conflicting SNP. If truly homozygous SNPs are listed as heterozygous, TReCASE will take them at face value while RASQUAL again will try to correct them. Because RASQUAL needs to use multiple SNPs to correct a wrong one, we only consider a scenario when splitting counts to 8 SNPs. Since most genes have less than 8 heterozygous SNPs, these simulation results represent an uncommon situation that favor the genotyping error correction mechanism of RASQUAL.

We consider the simulation setting when the over-dispersion parameters of negative binomial and beta-binomial are the same to match with the assumption made by RASQUAL. We illustrate the type I errors and powers for this simulation setup for a few values of over-dispersion parameters and several sample sizes (Figure 15). TReCASE controls type I error in all simulation setups. In contrast, RASQUAL controls type I error only if sample size is small and over-dispersion is small (Figure 15 (d)-(f)). The power of TReCASE remains similar as the proportion of genotyping errors increases, with some slight reduction of power when over-dispersion is large and eQTL effect size is large. Larger fraction of genotyping errors also reduces the power of RASQUAL in a magnitude slightly larger than for TReCASE (Figure 15 (a)-(c)).

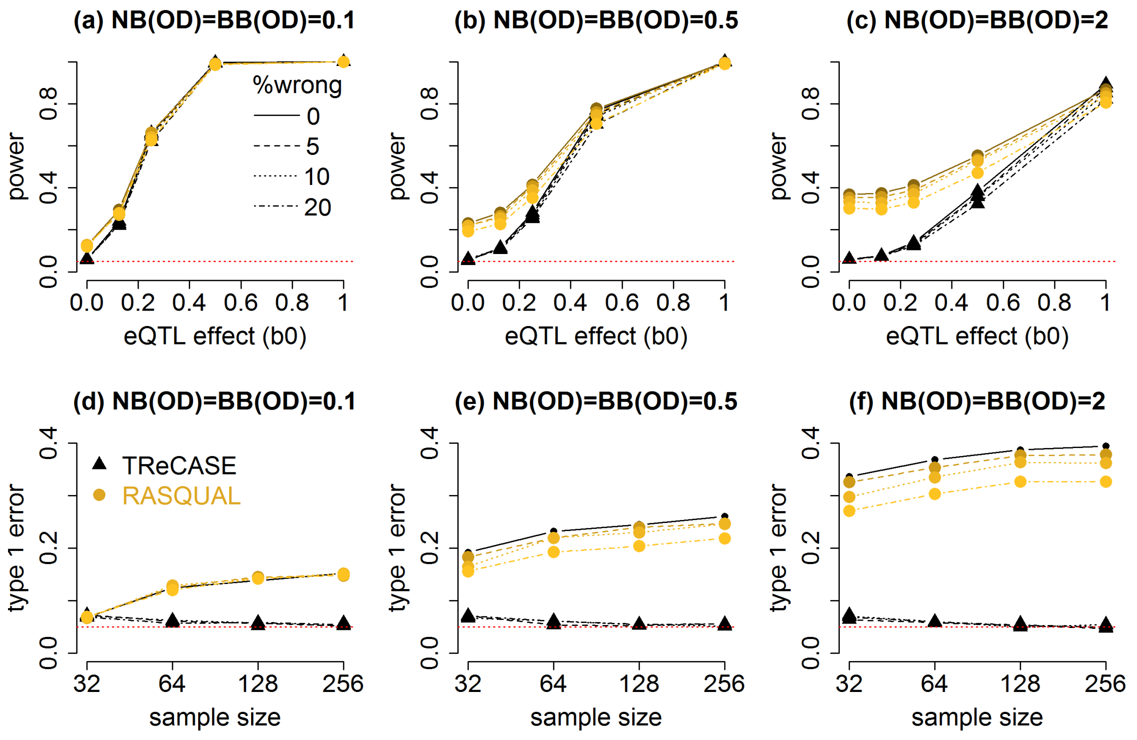

Figure 15: Type I errors and powers for a fitting TReCASE and RASQUAL with 0, 5%, 10%, or 20% of all the fSNPs being randomly flipped between homozygous and heterozygous. Sample size 64 was used.

##### C.5 Compare MatrixEQTL, TReCASE, and RASQUAL using Geuvadis dataset

###### C.5.1 Computational time

We performed eQTL mapping for all the genes for which there were at least 5 samples with at least 5 allele-specific reads. Since RASQUAL is very time consuming, we fit each gene in parallel. For timing comparability, we did the same for TReCASE. We limited total computational time per gene to be a week, and RASQUAL failed to finish within a week for 9 genes. We summarize the results for the remaining 14,427 genes. The average number of potential eQTL SNPs per gene was 2,000. The computational time of both TReCASE and RSQUAL increases nearly linearly with sample sizes, and TReCASE is more than 10 times faster than RASQUAL (Figure 16(a)). For the full dataset with sample size 280, if all the computation were done without any parallel computing, RASQUAL took 610 days and TReCASE took 53 days (Table 12).

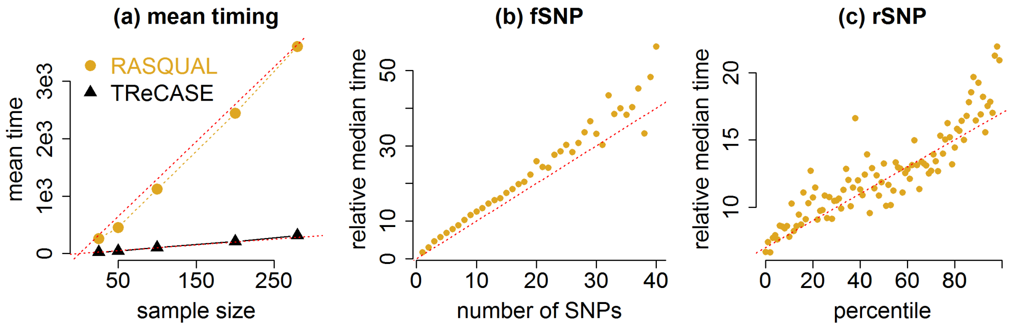

Figure 16: TReCASE vs RASQUAL timing for eQTL mapping. (a) The average computational time (seconds) for eQTL mapping per gene versus sample size. The dotted lines are $y = x$ and $y = 13x$. (b-c) The median of relative time for eQTL mapping per gene using RASQUAL versus TReCASE versus the number of feature SNPs (fSNPs) (with a reference line $y = x$) or the number of eQTL SNPs (rSNPs) (with a reference line $y = 7 + 0.1x$).

We also include the computational time for permutation p-value estimation by our method geoP, which uses a logistic regression to estimate the relation between nominal p-value and permutation p-value. Because the estimation of permutation p-value uses computationally very efficient linear regression implemented in MatrixEQTL (Shabalin, 2012), its computational time is comparable to one run of TReCASE. For 14,500 genes with 25, 50, 100, and 200 grid points per gene, it takes approximately 19, 23, 28, and 42 days, respectively (Table 12). Although it still takes substantial amount of time, it can be easily paralleled gene by gene.

Table 12: Comparison of computational time in days. 'one run': time for one round of eQTL mapping. 'est.perm.pval': time for estimation of permutation p-values using 100 grid points, or the range of computational time shown in the parenthesis using 25 to 200 grid points. 'total time': the total time needed for eQTL mapping and estimation of permutation p-values. The last row of the table shows the computational time of a modified version of TReCASE (Hu et al., 2015) where a score test is developed that improves computational efficiency. The TReCASE(score) method does not separate the initial run and permutation p-value calculation and thus we record the total time using 5,000 permutations.

| method | one run | est. perm.pval | total time |
|---|---|---|---|
| RASQUAL | 610 | 28 (19-42) | 638 (629-652) |
| TReCASE | 53 | 28 (19-42) | 81 (72-95) |
| TReCASE (score) | - | | 750 (750) |

We have developed a modified version of TReCASE to perform testing using score test (Hu et al., 2015). This TReCASE (score) method is computationally more efficient to perform permutations since some elements of the score test statistic can be calculated only once and used for many permutations. For 280 samples in the Geuvadis dataset, TReCASE (score) took 750 days for 5,000 permutations (Table 12), and thus it is doable within a week using 100+ computing jobs. However, one limitation of score test is that the p-values become less stable when sample size is relatively small, such as n=50 or 100. While the permutation p-values are still accurate, one should be cautious when using the nominal p-values with small sample sizes.

The relative computational time of RASQUAL versus TReCASE increases with respect to the number of fSNPs, i.e., the feature SNPs on which ASE is measured (Figure 16(b)) and the number of rSNPs , i.e., eQTL SNPs (Figure 16(c)). When genotype data were obtained by whole genome sequencing (e.g., GTEx dataset), there are about 10 times of fSNPs per gene (Figure 17) than the genotype data obtained from imputation (e.g., Geuvadis dataset). As a consequence, RASQUAL can be 100 times slower than TReCASE. For example, we compared the computational time of TReCASE and RASQUAL using GTEx v7 whole blood data, using a smaller local-eQTL window of 100kb flanking regions + gene body. Using a single CPU, TReCASE takes 14 days while RASQUAL takes 1,663 days. For this reason, we do not evaluate RASQUAL using the GTEx v8 data.

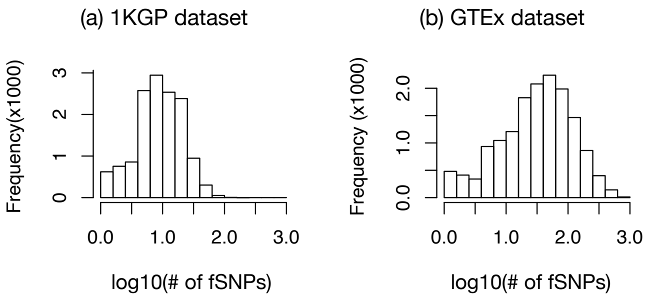

Figure 17: Number of fSNPs per gene. The distribution of the number of fSNPs per gene for (a) Geuvadis (1KGP) dataset or (b) GTEx dataset (whole blood, v7), respectively.

##### C.5.2 Choose a permutation p-value cutoff to control FDR

Given the permutation p-value for each gene, we can choose a permutation p-value cutoff to control FDR by evaluating q-value. We estimated the fraction of genes that follow the null distribution (denoted by $\pi_0$) by doubling the fraction of the genes with permutation p-values above 0.5, and then estimate FDR (or an approximation of q-value) for permutation p-value $\alpha$ by $\pi_0 \alpha N / D_\alpha$, where $N$ is the total number of genes and $D_\alpha$ is the number of genes with permutation p-value $\leq \alpha$. Since a larger proportion of genes has significant eQTLs, $\pi_0$ can be small and thus for some FDR cutoffs, the corresponding permutation p-value can be even larger than the FDR (Table 13). For example, to control FDR at 0.05, the p-value cutoffs are larger than 0.05. Here we use FDR cutoff 0.01 to choose permutation p-value cutoffs for the three methods.

Table 13: Permutation p-value cutoffs for different FDR cutoffs, using Geuvadis dataset with sample size 280.

| FDR | TReCASE | RASQUAL | MatrixEQTL |
|---:|---:|---:|---:|
| 0.001 | 0.001 | 0.0009 | 0.0005 |
| 0.01 | 0.014 | 0.011 | 0.009 |
| 0.05 | 0.092 | 0.071 | 0.074 |
| 0.1 | 0.211 | 0.161 | 0.189 |
| 0.2 | 0.474 | 0.362 | 0.477 |
| 0.25 | 0.616 | 0.470 | 0.631 |

##### C.5.3 Compare TReCASE vs. MatrixEQTL

As illustrated in Figure 2 of main text, TReCASE has much higher power than MatrixEQTL when sample size is relatively small (e.g., $n \leq 140$). However, such results do not answer a related but different question: how many of the eQTLs overlap and how do such overlaps change with respect to sample size. We first show that TReCASE vs. MatrixEQTL tend to identify the same set of eQTLs when sample size is large enough because the correlations of their p-values (in log scale) increases with sample size (Table 14).

Table 14: Correlations of TReC or TReCASE p-values vs MatrixEQTL p-values on -log10 scale

| Sample size (n) | TReC | TReCASE |
|---:|---:|---:|
| 35 | 0.72 | 0.67 |
| 70 | 0.89 | 0.77 |
| 140 | 0.95 | 0.85 |
| 280 | 0.97 | 0.87 |

Next, we compare the overlaps of the eQTL findings across sample sizes using q-value 0.01 as cutoff. Because we have more confidence on the eQTLs identified with larger sample sizes, we made three comparisons where each time, we compared the results of all the methods at all sample sizes versus the eQTL findings by one method using the maximum sample size of 280. TReCASE can identify most eGenes reported by MatrixEQTL (i.e., 95% when n=280) while MatrixEQTL still misses 42% of the eGenes identified by TReCASE when n=280 (Table 15 and 17). Since both MatrixEQTL and TReC use total expression but not ASE, as expected there is higher consistency when comparing MatrixEQTL vs. TReC than comparing MatrixEQTL vs. TReCASE (Table 16). It is also clear that sample size 35 is too small for reliable eQTL mapping. MatrixEQTL report no findings when n=35 and 19% of TReCASE eQTL findings cannot be found when sample size is 280.

Table 15: Comparison to the eGenes identified by TReCASE when sample size (n) is 280. The columns of "Recovered eGenes" are the proportion of TReCASE eGenes (when n=280) recovered by each method at each sample size. The columns of "eGenes missed by TReCASE" are the proportion of eGenes identified by each method (at each sample size) but missed by TReCASE when n=280.

| | Recovered eGenes | | | eGenes missed by TReCASE | | |
|---|---|---|---|---|---|---|
| n | MatrixEQTL | TReC | TReCASE | MatrixEQTL | TReC | TReCASE |
| 35 | 0 | 0.02 | 0.05 | - | 0.26 | 0.19 |
| 70 | 0.03 | 0.06 | 0.14 | 0.14 | 0.09 | 0.08 |
| 140 | 0.19 | 0.24 | 0.45 | 0.06 | 0.05 | 0.06 |
| 280 | 0.58 | 0.67 | 1 | 0.05 | 0 | 0 |

Table 16: Comparison to the eGenes identified by TReC when n=280.

| | Recovered eGenes | | | eGenes missed by TReC | | |
|---|---|---|---|---|---|---|
| n | MatrixEQTL | TReC | TReCASE | MatrixEQTL | TReC | TReCASE |
| 35 | 0 | 0.03 | 0.07 | - | 0.36 | 0.28 |
| 70 | 0.04 | 0.08 | 0.18 | 0.15 | 0.11 | 0.17 |
| 140 | 0.27 | 0.34 | 0.54 | 0.08 | 0.08 | 0.23 |
| 280 | 0.81 | 1 | 1 | 0.09 | 0 | 0.33 |

Table 17: Comparison to the eGenes identified by MatrixEQTL when n=280.

| | Recovered eGenes | | | eGenes missed by MatrixEQTL | | |
|---|---|---|---|---|---|---|
| samples | MatrixEQTL | TReC | TReCASE | MatrixEQTL | TReC | TReCASE |
| 35 | 0 | 0.03 | 0.07 | - | 0.37 | 0.28 |
| 70 | 0.05 | 0.09 | 0.2 | 0.09 | 0.13 | 0.2 |
| 140 | 0.31 | 0.37 | 0.57 | 0.07 | 0.12 | 0.28 |
| 280 | 1 | 0.91 | 0.95 | 0 | 0.19 | 0.42 |

##### C.5.4 Genomic locations of eQTLs

For each method, we selected the eGenes with q-value $\leq 0.01$ and for each eGene, we selected the SNP with the most significant p-value. Then we summarized the location of those eQTLs with respect to the corresponding genes. We observe enrichment around transcription starting site regardless of the strand of each gene (Figure 18). There is also a relatively weaker enrichment around transcription ending site (TES). Note that few eQTLs are located more than 10kb away from the corresponding gene. We also tried several cutoffs and noticed that when we select eQTLs with smaller p-value and thus reducing fraction of false positive discoveries, the fraction of eQTLs farther away from gene goes down.

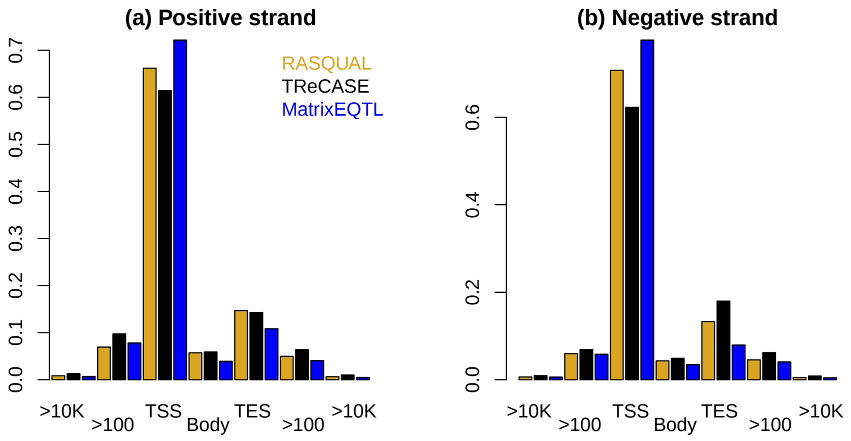

Figure 18: The distribution of the most significant eQTL for each eGene (q-value $\leq 0.01$). The region "TSS" or "TES" are defined as the region of 200bp (i.e., 100 bp on either side) around transcription starting site (TSS) or transcription ending site (TSS), respectively. The regions labeled ">100" are between 100 and 10kb away from TSS or TES.

##### C.5.5 Compare the results of RASQUAL vs. TReCASE

RASQUAL estimates one over-dispersion that is shared by its negative binomial and beta-binomial components and TReCASE estimates the over-dispersion parameters for these two components separately. We observe a quite clear pattern that the over-dispersion estimate from RASQUAL is very similar to the over-dispersion estimate of TReCASE negative binomial (TReC) component, but they are often much larger than the over-dispersion estimate of TReCASE beta binomial (ASE) component (Figure 19).

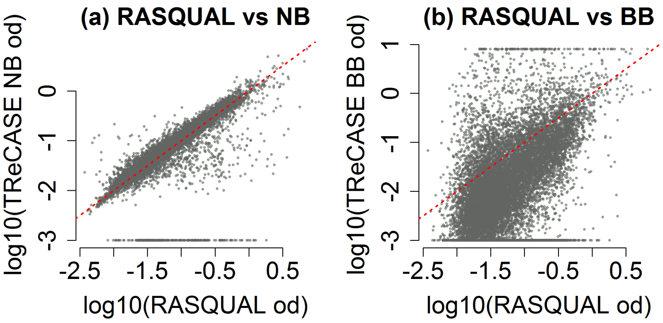

Figure 19: Comparing estimates of over-dispersion by RASQUAL or TReCASE. (a) TReCASE negative-binomial over-dispersion estimates and (b) TReCASE beta-binomial over-dispersion estimates. We trimmed log10(over-dispersion values) from TReCASE output to $[-3, 1]$ range.

We classified the fitted genes by their significance according to each of two methods (Table 18). Although the results of the two methods are consistent for the majority of the genes, there is some notable discrepancy for a subset of genes.

Table 18: Number of genes passing corresponding cutoff of q-values applied on permuted p-values.

| TReCASE | $[0.01, 1]$ | $[10^{-3}, 0.01)$ | $[10^{-4}, 10^{-3})$ | $[10^{-5}, 10^{-4})$ | $<$ 1e-5 | total |
|---|---|---|---|---|---|---|
| RASQUAL | | | | | | |
| $[0.01, 1]$ | 5463 | 1544 | 520 | 163 | 140 | 7830 |
| $[10^{-3}, 0.01)$ | 363 | 441 | 366 | 174 | 192 | 1536 |
| $[10^{-4}, 10^{-5})$ | 148 | 161 | 215 | 161 | 251 | 936 |
| $[10^{-5}, 10^{-4})$ | 78 | 100 | 112 | 91 | 242 | 623 |
| $<$ 1e-5 | 323 | 217 | 243 | 270 | 2449 | 3502 |
| total | 6375 | 2463 | 1456 | 859 | 3274 | 14427 |

To identify the potential sources for discrepancies, we conducted the following analyses. For each gene, we took the smallest p-value across multiple rSNPs by TReCASE and RASQUAL, truncated them at $10^{-15}$, and refer them as TReCASE p-value and RASQUAL p-value, respectively. We sought to explore the discrepancies of TReCASE and RASQUAL p-values by a linear regression with

$$y = \log_{10}(\text{TReCASE p-value}) - \log_{10}(\text{RASQUAL p-value})$$

as the response variable and 16 covariates (Tables 19-20):

- alternative allele frequency
- $\log_{10}$ p-value of $\chi^2$ test for Hardy Weinberg equilibrium
- estimated mapping error (Delta), which is an output of RASQUAL
- reference allele bias (Phi Bias), which is an output of RASQUAL
- the number of feature SNPs per gene, centered to median.
- the number of rSNPs per gene, on log scale.
- over-dispersion from total read counts ($\text{OD}_{NB}$) estimated by TReCASE, in log scale and centered.
- over-dispersion from allele-specific counts ($\text{OD}_{BB}$) estimated by TReCASE, in log scale and centered.
- the total allele-specific counts by RASQUAL, in log scale and centered.
- the total allele-specific counts by TReCASE, in log scale and centered.
- interaction of the previous two counts. Two methods count allele-specific reads differently. TReCASE count them at gene level while RASQUAL count them SNP by SNP. Therefore, if one read overlaps with two fSNPs, it will be counted twice. The potential degree of over-counting is illustrated at Figure 21.

- interactions of three covariates: RASQUAL allele-specific counts, the number of fSNPs, and beta binomial over-dispersion. Based on our simulations and study of information matrix we believe that p-value inflation of RASQUAL is caused by its model of ASReC. These three covariates are all important for the ASReC model.

- median p-value of RASQUAL across all rSNPs of a gene, using permuted genotype. This quantity measures the magnitude of RASQUAL type I error for this gene.

All covariates were normalized to have standard deviation of 1.

Table 19: Type 1 (sequential) and Type 3 (added last) ANOVAs for linear regression analysis of $\log_{10}$(TReCASE p-value) - $\log_{10}$(RASQUAL p-value). The direction (Dir.) indicates whether RASQUAL (R) or TReCASE(T) has smaller p-value.

| Parameter | Dir. | Type 1 $R^2$ | P-val | Type 3 $R^2$ | P-val | Marg. $R^2$ |
|---|---|---|---|---|---|---|
| $\mathrm{OD}_{BB}$ | R | 6.7 | 6e-244 | 8.2 | 3e-294 | 6.7 |
| $\mathrm{ASReC}_R$ | R | 2.7 | 8e-101 | 1 | 2e-39 | 0.6 |
| n-fSNP | R | 1.9 | 1e-71 | 1.6 | 4e-63 | 2.7 |
| $\mathrm{ASReC}_R$ :n-fSNP | R | 0.5 | 2e-19 | 0.7 | 4e-26 | 0 |
| $\mathrm{ASReC}_R$ :$\mathrm{OD}_{BB}$ | R | 5 | 1e-185 | 3.2 | 1e-118 | 1.3 |
| n-fSNP:$\mathrm{OD}_{BB}$ | R | 0.8 | 6e-31 | 1 | 1e-37 | 0.4 |
| $\mathrm{ASReC}_R$ :n-fSNP:$OD_{BB}$ | R | 0.2 | 4e-8 | 0.2 | 2e-10 | 0.1 |
| n-rSNP | T | 0 | 0.73 | 0 | 0.52 | 1 |
| $\mathrm{OD}_{NB}$ | T | 0 | 0.009 | 0 | 0.004 | 0.8 |
| $\mathrm{ASReC}_T$ | T | 0.4 | 1e-15 | 0.4 | 3e-16 | 0.1 |
| $\mathrm{ASReC}_R$ :$\mathrm{ASReC}_T$ | R | 0.3 | 6e-13 | 0.2 | 1e-10 | 0 |
| AF | T | 0 | 0.004 | 0.1 | 0.002 | 0 |
| HWE $\chi^2$ | T | 0.2 | 5e-9 | 0.2 | 1e-8 | 0.3 |
| Mapping error | T | 0 | 0.09 | 0 | 0.24 | 1.1 |
| Ref. Allel Bias | R | 0.3 | 2e-13 | 0.3 | 2e-13 | 2.1 |
| Med(perm-p) | T | 0 | 0.97 | 0 | 0.97 | 0.4 |

This linear model explains 15% of the variance of $y$. More variance of $y$ can be explained by this model if we only consider a subset of genes with more discrepant p-values. For example, for a subset of genes passing a cutoff of $|y| \geq 5$ or $|y| \geq 10$,

Table 20: Linear regression of $y = \log_{10}$(TReCASE p-value) - $\log_{10}$(RASQUAL p-value) versus a set of potential factors. $OD_{BB}$ and $OD_{NB}$ indicate $\log_{10}$ over-dispersion for beta-binomial and negative binomial, respectively. n-fSNP and n-rSNP indicate the number of feature SNPs and regulatory SNPs, respectively. Subscript $_R$ and $_T$ indicate RASQUAL and TReCASE, respectively.

| Parameter | Est | SE | P-val | Marg.Est | Marg P-val |
|---|---|---|---|---|---|
| intercept | 0.18 | 0.14 | 0.2 | 0.31 | 1.6e-43 |
| $OD_{BB}$ | 1.04 | 0.03 | 3e-294 | 0.68 | 3e-213 |
| $ASReC_R$ | 0.81 | 0.06 | 2e-39 | 0.20 | 4e-19 |
| n-fSNP | 0.44 | 0.03 | 4e-63 | 0.43 | 4e-84 |
| $ASReC_R$ :n-fSNP | 0.24 | 0.023 | 4e-26 | 0.013 | 0.51 |
| $ASReC_R$ :$OD_{BB}$ | 0.52 | 0.022 | 1e-118 | 0.26 | 7e-42 |
| n-fSNP:$OD_{BB}$ | 0.29 | 0.022 | 1e-37 | 0.13 | 2e-12 |
| $ASReC_R$ :n-fSNP:$OD_{BB}$ | 0.11 | 0.017 | 2e-10 | -0.04 | 0.0027 |
| n-rSNP | -0.014 | 0.022 | 0.52 | 0.26 | 6e-32 |
| $OD_{NB}$ | -0.07 | 0.024 | 0.004 | 0.24 | 1e-26 |
| $ASReC_T$ | -0.46 | 0.056 | 3e-16 | 0.084 | 2e-4 |
| $ASReC_R$ :$ASReC_T$ | 0.12 | 0.019 | 1e-10 | 0.037 | 0.048 |
| AF | -0.061 | 0.02 | 0.002 | -0.044 | 0.046 |
| HWE $\chi^2$ | -0.12 | 0.02 | 1e-8 | -0.16 | 3e-12 |
| Mapping Error | -0.028 | 0.024 | 0.24 | 0.28 | 2e-36 |
| Ref. Allel Bias | 0.19 | 0.026 | 2e-13 | 0.38 | 6e-65 |
| Med(perm p) | -0.001 | 0.034 | 0.97 | -0.16 | 5e-13 |

this linear model explains 42% or 57% of the variance of $y$, respectively. We note that the more discrepant set of genes we select, the higher fraction of genes with smaller p-values by RASQUAL (Figure 20 (a)), suggesting stronger discrepancies are more likely due to the inflation of Type I error by RASQUAL.

We observe that three factors, beta-binomial over-dispersion ($OD_{BB}$), the number of fSNPs (n-fSNP), and RASQUAL style ASReC ($ASReC_R$), along with interactions of these terms have strongest associations with the discrepancy of the two methods (Table 20). Larger values of these three factors are all associated with smaller RASQUAL p-values. This is consistent with our findings that the beta-binomial component of RASQUAL treats multiple fSNPs within a sample as independently

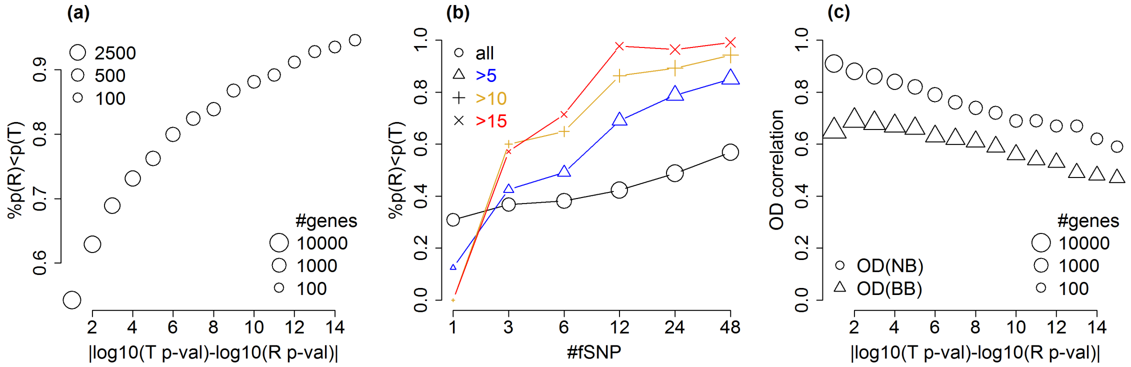

Figure 20: Comparison of TReCASE and RASQUAL results using Geuvadis dataset. We use "T p-val" and "R p-val" as abbreviations of "TReCASE p-value" and "RASQUAL p-value", respectively. "%p(R) < p(T)" denotes the proportion of genes with RASQUAL p-value smaller than TReCASE p-value. (a) Genes with more discrepant p-values tend to smaller RASQUAL p-values. (b) The genes with larger number of fSNPs also tend to have smaller RASQUAL p-values. Different point symbols indicate the genes with the absolute value of the difference between $\log_{10}$(TReCASE p-value) and $\log_{10}$(RASQUAL p-value) is larger than certain threshold. (c) When there are larger discrepancies of p-values, the over-dispersion estimates by RASQUAL are less similar to either negative binomial (NB) or beta-binomial (BB) over-dispersion estimates by TReCASE.

distributed, which causes larger inflation of type I error. In addition, more discrepant genes also tend to have weaker correlations of over-dispersion estimates between TReCASE and RASQUAL (Figure 20 (c)).

Among other covariates, the following relations are notable. Significant associations with reference allele bias and Hardy-Weinberg disequilibrium suggest the advantage of RASQUAL to model these factors. Smaller TReCASE p-values are associated with the case when we observed relatively higher TReCASE style allele-specific counts ($\text{ASReC}_T$), which suggests that the ASReC by RASQUAL and TReCASE are different, most likely due to double counting by RASQUAL (Figure 21).

Median RASQUAL p-value using permuted data (Med(perm p)) is very signifi-

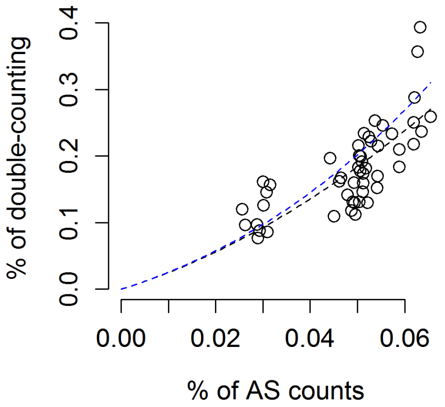

Figure 21: Estimating double-count in real data

We again used 30 samples from PRJNA385599. For each sample we counted number of allele-specific reads using our TReCASE procedure and produced fraction with respect to total number of reads (x) and will plot on x scale. In addition, for the reads overlapping several heterozygous SNPs we counted such read several time - once for each SNP (define this number as z) Z is inflated with overcounting. We quantify this excess of counts by defining $y = z/x - 1$ and plotting them on the y axis. 10 of the samples in this dataset were measured with both 150bp reads and shorter 75bp reads. They are plotted separately with 10 points around 3% allele-specific counts representing summary for shorter reads.

cant in marginal model, but becomes much less significant in the joint model. This is expected because inflation of type I error is also associated with other factors in the join model (see Section C.4.4 for more details). In both cases, smaller RASQUAL p-value using permuted data are associated with smaller RASQUAL p-values.

We further examine the discrepancy of significant findings by RASQUAL and TReCASE with respect to the number of fSNPs. We classified the genes to be significant or not at several FDR cutoffs and plotted them versus the number of fSNPs. The fraction of significant findings of both methods generally grows with respect to the number of fSNPs (Figure 22(a) and (d)), which is expected since the number of allele-specific reads would also be higher with more fSNPs. However, this fraction grows quicker for RASQUAL than TReCASE, and it increases regardless the significance level of TReCASE (Figure 22(b) and (c)). This suggests that the association between the number of fSNPs and RASQUAL p-values may not depend on the actual strength of eQTL signals and thus implies inflated type I error. In contrast, conditioning on being significant or insignificant using RASQUAL method we see much weaker association between the number of fSNPs and the fraction of significant TReCASE findings (Figure 22(e) and (f)). In fact, given significant RASQUAL results, the number of significant TReCASE findings has slight decrease as number of fSNPs increases (Figure 22(e)). This is likely because RASQUAL tends to find higher fraction of false positive when the number of fSNPs is large.

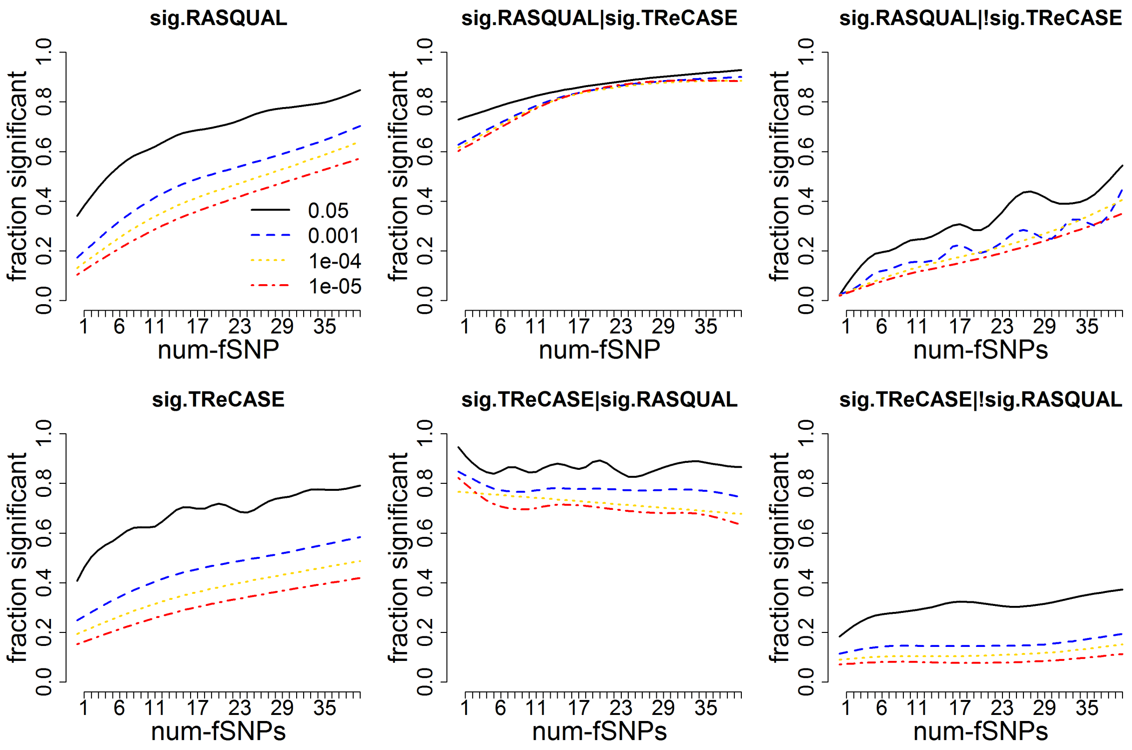

Figure 22: Method discrepancy conditioned on significance status of each method. We classify the genes into significant or not significant category using FDR cutoffs presented in the legend: 0.05, 1e-3, 1e-4 and 1e-5 plotted vs number of fSNPs. The curve is obtained using a spline. Panels (a) and (d) consider overall dependency of fraction of genes found to be significant plotted versus number of fSNPs. Panel (b) considers proportion of genes passing a cutoff in RASQUAL model for all the genes passing cutoff for TReCASE. Panel (e) does it other way around - fraction of significant genes found by TReCASE among the genes significant in RASQUAL. Panels (c) and (f) provide similar curves for fraction of genes found to be significant by one of the methods, given that they weren't found to be significant by the other method.

##### C.5.6 eQTL mapping using permuted genotype data

In this subsection, we evaluate potential type I error inflation of TReCASE and RASQUAL using permuted genotype data. We only include the potential *cis*-acting eQTLs in our evaluation because ASE is only informative for *cis*-eQTL mapping. To identify *cis*-acting eQTLs, we test whether the eQTL effects estimated by TReC and ASE are the same by a cis-trans test (Sun, 2012), and consider the cases with cis-trans test p-value $> 0.05$ as the potential *cis*-acting eQTLs. For TReCASE method, we consider the standard TReCASE using likelihood ratio test (LRT) (Sun, 2012) as well as another version using score test (Hu et al., 2015).

From the distribution of all the eQTL p-values, it is clear that RASQUAL has severely inflated type I error (Figure 23(c,f)). This is consistent with our simulation results (Section C.4.4) and comparison of RASQUAL and TReCASE using the Geuvadis dataset (Section C.5.5). With sample size of 280, the p-value distribution from TReCASE (score) is slightly deviated from uniform distribution (Figure 23(a,d)), though such deviation becomes larger for smaller sample size of 100 (Figure 24(a,d)). Therefore, when using TReCASE (score) method for small sample size, we recommend using permutation p-values rather than the p-values from asymptotic distribution.

We observed that TReCASE also has slight inflated type I error. This may be due to model mis-specification for some genes, either because the distribution assumption is not accurate or missing some covariates (e.g., genetic effect are missing after permuting genotype data). Such inflation of type I error can be removed after trimming outlier values using an approach implemented in DESeq2 (Love et al., 2014). An observation is defined as an outlier if it Cook's distance is larger than a thresholds, and found the threshold of $4/n$ (Hardin et al., 2007) effectively removes the inflation of type I error (Figure 25).

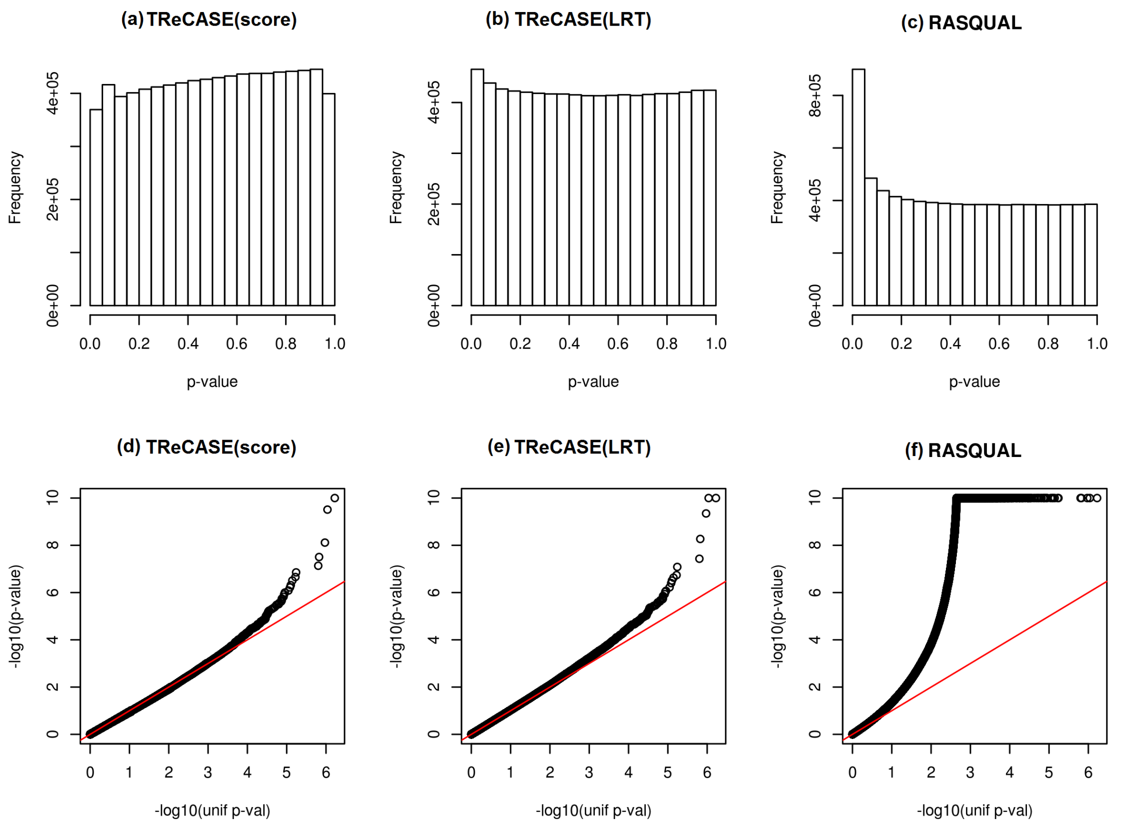

Figure 23: The distributions ((a)-(c)) and QQ-plots ((d)-(f)) of eQTL p-values using permuted genotypes by three methods: TReCASE (LRT), TReCASE(Score) and RASQUAL, using Geuvadis dataset with sample size 280.

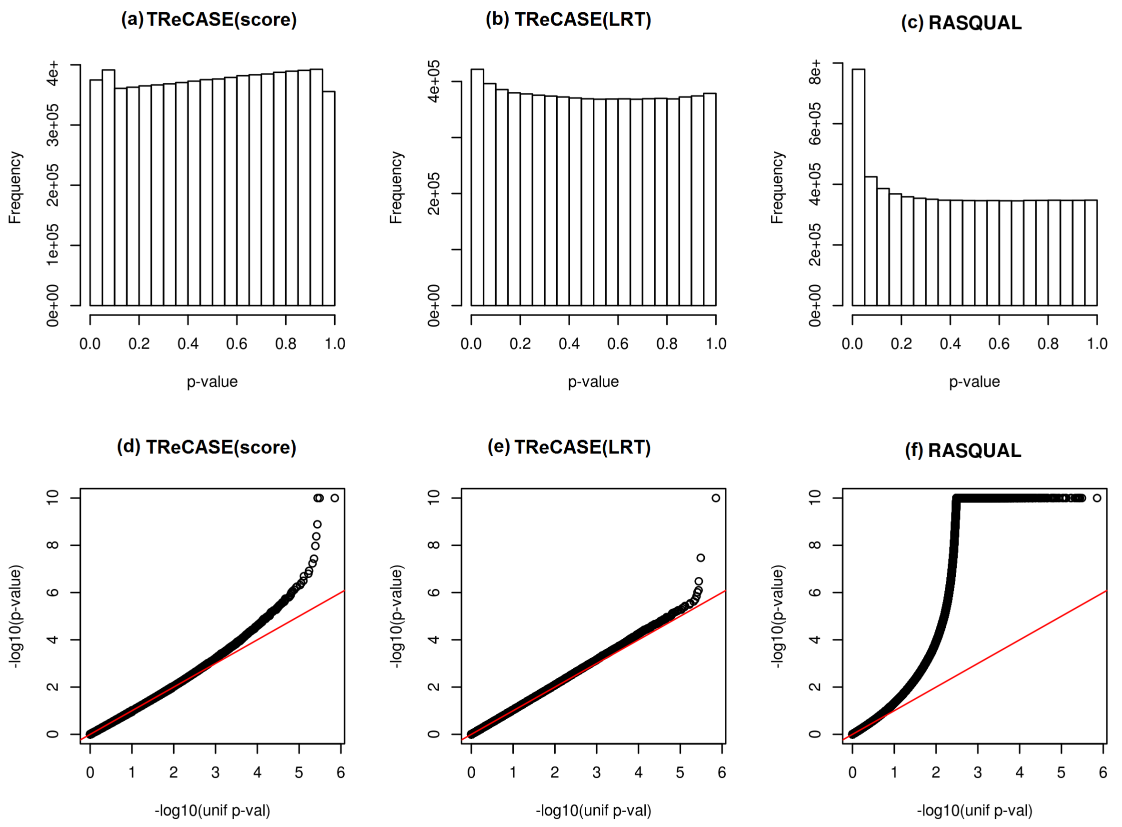

Figure 24: The distributions ((a)-(c)) and QQ-plots ((d)-(f)) of eQTL p-values using permuted genotypes by three methods: TReCASE (LRT), TReCASE(Score) and RASQUAL, using Geuvadis dataset with sample size 100.

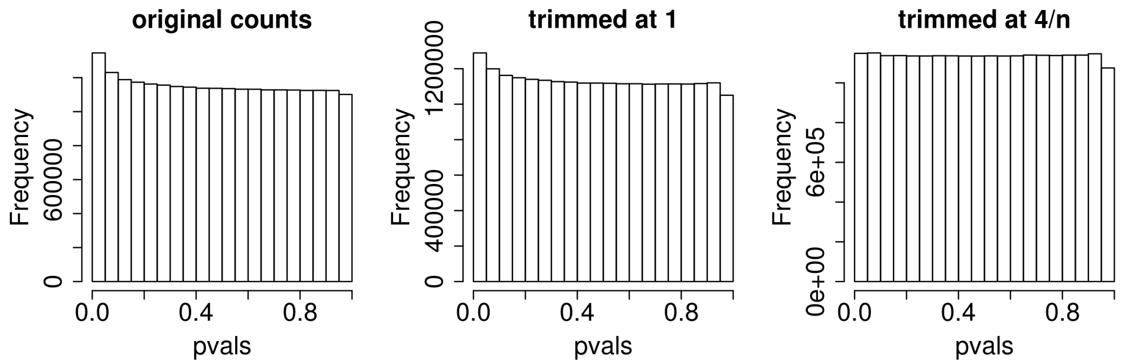

Figure 25: Fitting the data with permuted genotypes using TReC model after trimming counts with significant Cook's distances

#### C.6 Compare MatrixEQTL vs. TReCASE using Geuvadis and GTEx v7 data

As mentioned earlier in Section C.5.1, it is computationally infeasible to run RASQUAL on GTEx data. Due to the large number of feature SNPs in GTEx data, RASQUAL can be 100 times slower than TReCASE. Here we make a systematic comparison of TReCASE versus MatrixEQTL using both Geuvadis and GTEx v7 data.

In Geuvadis dataset with permutation p-values estimated by geoP, MatrixEQTL was able to identify 3,336 genes at $q = 0.01$ cutoff versus 6,740 found by TReCASE (Table 21). If the permutation p-values are estimated using eigenMT, MatrixEQTL results identify only 2,773 genes to be significant at q-value 0.01. This is because eigenMT is more conservative than geoP for the permutation p-values that correspond to q-value of 0.01. The estimated proportion of nulls using gene-specific permutation p-values are 23.5%, 41.5%, and 94.2% for TReCASE (geoP) , MatrixEQTL (geoP), and MatrixEQTL (eigenMT), respectively.

Table 21: Number of genes passing corresponding q-value cutoffs (that were estimated using permuted p-values estimated by geoP) in Geuvadis dataset by TReCASE vs MatrixEQTL.

| MatrixEQTL<br>TReCASE | $[0.01, 1]$ | $[10^{-3}, 0.01)$ | $[10^{-4}, 10^{-3})$ | $[10^{-5}, 10^{-4})$ | $<$ 1e-5 | total |
|---|---|---|---|---|---|---|
| $[0.01, 1]$ | 7543 | 167 | 39 | 16 | 41 | 7806 |
| $[10^{-3}, 0.01)$ | 1727 | 223 | 61 | 26 | 26 | 2063 |
| $[10^{-4}, 10^{-5})$ | 666 | 147 | 70 | 35 | 55 | 973 |
| $[10^{-5}, 10^{-4})$ | 416 | 104 | 72 | 42 | 88 | 722 |
| $<$ 1e-5 | 858 | 298 | 241 | 223 | 1362 | 2982 |
| total | 11210 | 939 | 483 | 342 | 1572 | 14546 |

GTEx dataset (v7, whole blood) shows more dramatic gain from MatrixEQTL to TReCASE (Table 22) with only 1,878 genes passing $q = 0.01$ cutoff for MatrixEQTL + geoP versus 7,850 found by TReCASE + geoP. Applying eigenMT to MatrixEQTL results would make the results even more conservative declaring only 1,662 genes to be significant at $q = 0.01$. The estimated proportion of nulls using gene-specific permutation p-values are 21.2%, 57.4%, and 100% for TReCASE (geoP), MatrixEQTL (geoP), and MatrixEQTL (eigenMT), respectively.

Table 22: Number of genes passing corresponding q-value cutoffs (that were estimated using permuted p-values estimated by geoP) in GTEx dataset (v7 whole blood) by TReCASE vs MatrixEQTL.

| MatrixEQTL | $[0.01, 1]$ | $[10^{-3}, 0.01)$ | $[10^{-4}, 10^{-3})$ | $[10^{-5}, 10^{-4})$ | $<$ 1e-5 | total |
|---|---|---|---|---|---|---|
| TReCASE | | | | | | |
| $[0.01, 1]$ | 8721 | 36 | 11 | 2 | 7 | 8777 |
| $[10^{-3}, 0.01)$ | 2067 | 71 | 26 | 11 | 9 | 2184 |
| $[10^{-4}, 10^{-5})$ | 927 | 78 | 28 | 15 | 11 | 1059 |
| $[10^{-5}, 10^{-4})$ | 543 | 58 | 30 | 20 | 28 | 679 |
| $<$ 1e-5 | 2491 | 264 | 212 | 152 | 809 | 3928 |
| total | 14749 | 507 | 307 | 200 | 864 | 16627 |

Next, we study whether the eSNPs (the most significant eSNPs per gene) found by MatrixEQTL and TReCASE are located in different genomic regions, in terms of the 18 chromatin states classification provided by Roadmap Epigenomic Consortium (Kundaje et al., 2015). We consider the results of the two methods are concordant if their eSNPs of the same gene are within 10kb. Considering the SNPs significant at permutation p-value $\alpha = 0.01$ level and contrasting them to the genes non-significant (at $\alpha = 0.1$ level). Then each gene can be assigned to one of 8 categories based on 3 factors, whether the eSNPs found by MatrixEQTL and TReCASE are concordant and whether the eQTL association is significant for each method. We observed the distribution of these 8 groups has large difference for a few chromatin states (Figure 26). The eQTLs identified by both methods or by TReCASE only are less likely located in the Weak Repressed PolyComb (ReprPCwk) or Quiescent/Low (Quies) regions. We observed similar patterns in the results from GTEx (v7 whole blood) dataset (Figure 27).

Alternatively, we considered a different classification from Roadmap Epigenomic Consortium (Kundaje et al., 2015) in which certain DNase enriched regions were classified as promoter or enhancer. For these genes we estimated the probability of the SNP falls into a promoter or enhancer region based on the significance level by each method and distance from the gene to the corresponding eSNP (Table 23). We observe that in both datasets stronger TReCASE p-value is associated with higher

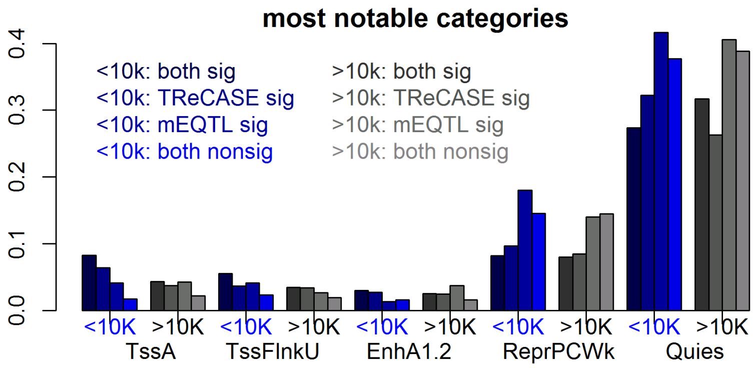

Figure 26: Distribution of eQTLs (from Geuvadis dataset) in different chromatin states. Here mEQTL stands for MatrixEQTL. For each of the 8 categories labeled in the top-left of the plot, we calculated the proportion of eQTLs located in each of the 18 chromatin states. Only the 5 chromatin states with larger difference across the 8 categories are shown. These groups are: (1) TssA - Active TSS, (2) TssFlnkU - Flanking TSS Upstream, (3) EnhA1 - Active Enhancer 1 and Active Enhancer 2, (4) ReprPCwk - Weak Repressed PolyComb and (5) Quies - Quiescent/Low

chance to be located within enhancer or particularly promoter while the signal for MatrixEQTL is weaker to non-existent. This observation suggests that incorporation of ASReC not only increases power, but also improves precision.

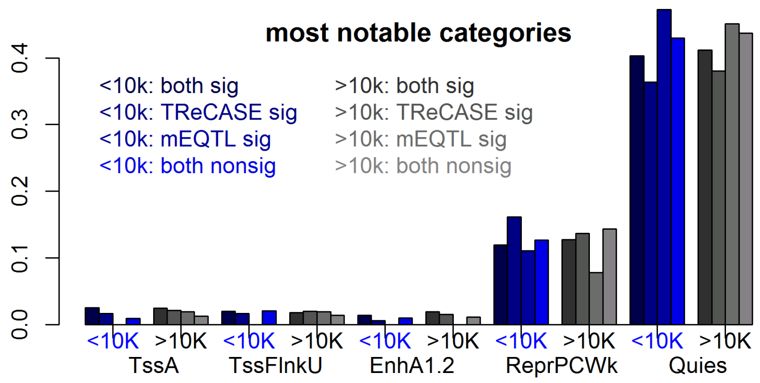

Figure 27: Distribution of eQTLs (from GTEx v7 whole blood dataset) in different chromatin states. The categories are the same as in the previous Figure.

Table 23: Association between eQTL signals and promoter or enhancer status. We fit a logistic regression with promoter/enhance status as response variable and with eQTL p-values and distance between the gene and the corresponding eSNP as predictors. The eQTL p-values are on negative $\log_{10}$ scale, and distance is on $\log_{10}$ scale with 1 added. The distance from a gene to a SNP within this gene is considered to be 0.

| | Geuvadis | | | | GTEx v7 whole blood | | | |
|---|---|---|---|---|---|---|---|---|
| | promoter | | enhancer | | promoter | | enhancer | |
| | coef | p-val | coef | p-val | coef | p-val | coef | p-val |
| Intercept | -2.5 | 0 | -3.21 | 8e-290 | -3.7 | 4e-254 | -3.6 | 8e-253 |
| $p_{mEQTL}$ | 0.01 | 0.37 | -0.02 | 0.07 | -5e-5 | 0.99 | -0.02 | 0.07 |
| $p_{TReCASE}$ | 0.05 | 8e-5 | 0.02 | 0.001 | 0.011 | 0.03 | 0.01 | 0.0003 |
| distance | -0.06 | 1e-15 | -0.005 | 0.58 | -0.014 | 0.18 | 0.003 | 0.76 |
